# Domain Insertion Permissibility is a Measure of Engineerable Allostery in Ion Channels

**DOI:** 10.1101/334672

**Authors:** Willow Coyote-Maestas, Yungui He, Chad L. Myers, Daniel Schmidt

## Abstract

Allostery is a fundamental principle of protein regulation that remains poorly understood and hard to engineer, in particular in ion channels. Here we use human Inward Rectifier K^+^ Channel Kir2.1 to establish domain insertion ‘permissibility’ as a new experimental paradigm to identify engineerable allosteric sites. We find that permissibility is best explained by dynamic protein properties, such as conformational flexibility. Many allosterically regulated sites in Kir2.1 or sites equivalent to those regulated in homologs, such as G-protein-gated inward rectifier K^+^ channels (GIRK), have *differential* permissibility; that is, for these sites permissibility depends on the structural properties of the inserted domain. Our data and the well-established link between protein dynamics and allostery led us to propose that differential permissibility is a metric of both existing and latent allostery in Kir2.1. In support of this notion, inserting light-switchable domains into either existing or latent allosteric sites, but not elsewhere, renders Kir2.1 activity sensitive to light.

## Introduction

Allostery is the phenomenon in proteins where the state of proximal sites (e.g., ligand binding to a sensor domain) is coupled to the state of distal sites (e.g., the activity of an effector domain). In nature, allosterically regulation arises from random recombination of functionally and structurally discrete protein domains or motifs. For example, many plant photoreceptors exhibit a highly modular architecture comprised of domains that provide a different function to the receptor^1^. In the lab, we recombine domains or motifs to generate synthetic proteins. For example, antibodies are joined end-to-end with signaling domains to create chimeric T-cell receptors for immunotherapy^2^. In both nature and the laboratory, how these components become coupled is essentially trial-and-error. That blind trial and error can, through natural selection over billions of years, progressively lead to optimized design was Darwin's groundbreaking concept for the evolution of natural systems^3^. In the lab, however, we need to accomplish this task in less time and with greater efficiency.

There have been, of course, several pioneering studies to rationally engineer allostery in proteins. Some rely on ‘manual’ recombination of domains with desired properties, either end-to-end or by in-frame domain insertion^4–9^. Unfortunately, this process requires tedious optimization of domain placement, and any successes can only be rationalized post hoc. Other methods that have guided the study and engineering of allostery in soluble proteins, such as random circular permutation^10^, NMR^11^, and computational approaches^12–14^, rely heavily on structural information or statistical methods. Thus, they remain relatively inaccessible to membrane proteins, for which there are fewer structures and homologs.

One class of membrane proteins that are particularly challenging to rationally engineer are ion channels. Ion channels play critical roles in the biological signaling processes that determine the operation of cells and networks of the brain and the heart and are thus major drug targets^15^. Virtually every aspect of ion channel gating relies on allosteric regulation, and many drugs achieve their therapeutic effect through allosteric modulation^16^. Being able to engineer the allosteric regulation of ion channels *de novo,* for example as chemo-or optogenetic tools, would enable an unprecedented fine-tuned control and exploration of how individual channels contribute to cellular functions^17,18^.

Concepts and models of allostery that could aid us in this task have been continuously refined since the description of haemoglobin, the prototypical allosteric protein^19,20^. In fact, in recent years, driven by the discovery that intrinsically disordered proteins exhibit allosteric regulation, ‘ensemble’ models of allostery have supplemented ‘structure-based allostery’ models^21^. We argue that a unified theory of allostery, to be useful for protein engineering, must also account for how allostery emerges in both natural and synthetic proteins.

For allosteric regulation to evolve through natural selection, the allosteric modulator and the modulated protein must *simultaneously possess* compatible allosteric signaling pathways that impart a selectable advantage. The most parsimonious way this can happen is if the required signal transduction networks *already exist* within the modulated protein. The term latent allostery has been coined for such hidden –latent– phenotypes that are not under selection, but that could be exploited in the same protein if selection pressures changes^22–24^.

To see how these latent functions can arise, consider a (hypothetical) ancestral unregulated channel protein whose conformation space is described by a highly favored closed (ground) state and a barely populated open (excited) state **Fig. 1A**). Mutations acquired by neutral drift, or random domain recombination could alter this free energy landscape such that ground and excited state structures remain (i.e., their positions in conformational space), but their relative stability (free energy) changes (**Fig. 1B**). If the excited state becomes more stable, then at the population level this hypothetical channel can perform some selectable function (e.g., allosteric regulation). Furthermore, it can do so with reasonable kinetics if a favorable ‘path’ of defined intermediate states exists that is void of large activation barriers and that connects ground and excited states. Collectively, we refer to all amino acids involved in these conformational transitions as an ‘exploited’ allosteric signaling network.

**Figure 1:**
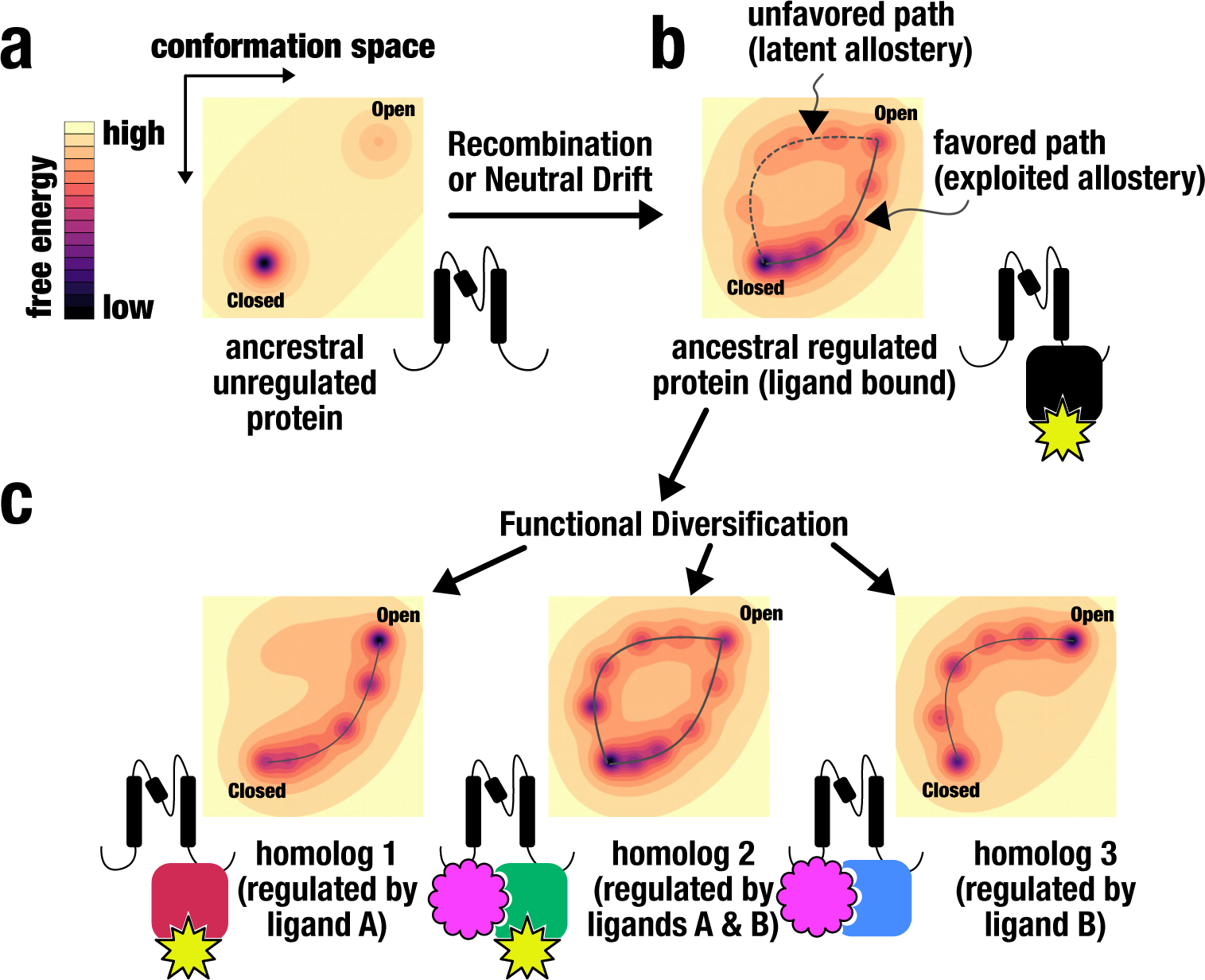
Latent and exploited allosteric regulation can be discovered by measuring domain insertion permissibility. **(a)** Energy landscape for a hypothetical unregulated ancestral protein. **(b)** Through domain recombination and/or neutral drift, and new energetic features are introduced into an ancestral protein that may or may not be exploited for allosteric regulation. **(c)** Subsequent diversification refines and optimize the features to give rise to homologous protein that can respond to different ligands.

While a specific function (e.g., allosteric regulation) is selected for when it provides an advantage under specific selection pressures, nothing limits the enabling mutations or domain recombination to provide *only* the selected function. It is entirely possible that other latent functions were introduced as byproducts (‘spandrels’^25^) that may or may not evolve into new phenotypic traits ^22^. From the viewpoint of the conformational landscape described above, a latent trait can be considered a set of alternative intermediate conformations that connect the ground and excited state, that nevertheless are less favorable perhaps due to large activation energy barriers and higher free energies. A protein that possesses such latent signal pathways can nevertheless utilize them to diversify function when selection pressures change. Therefore, this protein can adapt without major changes to protein architecture simply by acquiring a few mutations that cause of new allosteric modulator to bind, which in turn stabilizes the previously unfavorable signaling pathway (**Fig. 1C**).

Several studies have shown that latent phenotypes are co-opted in a number of different biological contexts, including soluble proteins such as enzymes^26^, hormone receptors^27^, and transcription factors^28^. As another example, a scaffold protein (Ste5) allosterically regulates Erk-like kinases that diverged *before* the evolution of Ste5 itself, implying that the allosteric potential to be regulated by Ste5 *was already present* at that point^24^.

How is latent allostery relevant to ion channels? The majority (43 out of 45) of human ion channel families appeared in the earliest metazoan^29,30^, so any subsequent functional diversification could conceivably be the result of leveraging latent regulatory mechanisms that existed in ancestral ion channel clades. However, it is unclear whether (1) ancestral channels used latent pathways to diversify, (2) modern ion channels still possess latent allosteric pathways, and (3) whether these pathways can be leveraged to engineer new allosteric regulation into channels. As a necessary first step to answering these questions, we here show how one can experimentally measure both exploited and latent allostery in ion channels in a way that does not rely on sequence or structural information. Instead, we measure how ‘permissive’ each ion channel site in is to insertion of domains with different biophysical properties (**Supp. Fig. 1**). We furthermore demonstrate this framework of measuring latent allostery in ion channels is useful to rationally engineer them to endow them with useful functions.

## Results

### A scalable, high-throughput domain insertion profiling pipeline for ion channels

Allosteric regulation requires that an allosteric site can discriminate and change in response to a perturbation and propagate information to change the state at a coupled distant site. Therefore, allosteric sites must be able to allow some mutations but be sensitive to the nature of that mutation. For these reasons, we hypothesized that sites with latent allosteric potential would have context-dependent mutability. To test this idea, we mimicked protein evolution and inserted protein domains (PDZ, Cib81, GSAG_2x_, GSAG_3x_) with different properties into nearly every amino acid position of Kir2.1 to measure site-specific domain insertion ‘permissibility’. Domain insertion ‘permissibility’ is defined as the site-specific ability of Kir2.1 to accept a domain insertion that does not disrupt folding, assembly, and trafficking to the cell surface. Kir2.1 is a good model for this proof-of-concept, because extensive functional studies are available including characterized mutations that are linked to the disruption of allosteric signaling and result in disease (e.g., Long QT syndrome^31^), high-resolution crystal structures that are available across phylogeny^32–36^, and the great allosteric regulatory diversity between homologs (e.g., G-proteins and ATP^37^). We used the 10kDa syntrophin PDZ domain (PDB 2PDZ) because it is well structured and has been used to study how large inserting large domains with known function disrupt recipient protein activity^38^. Cryptochrome-interacting basic-helix-loop-helix (Cib81) is a similarly sized domain that forms a two-component switchable system with its binding partner, CRY2, after blue-light illumination^39^. We included flexible GSAG_x2_ and GSAG_x3_ linkers to establish a permissibility baseline.

To generate insertion libraries, we used a saturation domain insertion profiling method, domain insertion profiling with sequencing (DIPSeq)^40^. DIPSeq uses a MuA transposase to insert an antibiotic cassette into random positions of a gene (i.e., all six reading frames) (**Supp. Fig. 1**). Upon antibiotic selection of variants with insertions, we replaced this cassette with a domain of interest (Cib81, PDZ, GSAG_x2_, and GSAG_x3_) using restriction sites at transposon ends that double as flexible linkers. To determine which amino acid positions are permissive and non-permissive to domain insertions, we leveraged the fact that for a channel to be expressed on the cell surface it must fold, oligomerize, and surface traffic^41^. Therefore, only permissive insertion variants can be fluorescently labeled via a FLAG epitope tag we inserted into an extracellular loop of Kir2.1 (S116) that does not interfere with channel function^42^. In this way, we collect two cell populations by fluorescence-activated cell sorting (FACS) (**Supp. Fig. 2**): those that express channel variants but don't surface express (EGFP^high^/anti-FLAG^low^) and those that do surface express (EGFP^high^/ anti-FLAG^high^). From both populations, we isolated and sequenced plasmid DNA, and aligned reads with the DIPSeq alignment pipeline^40^. We calculate permissibility as site-specific enrichment between 'not surface-expressed' to 'surface-expressed' insertion variants. Apart from some regions near the N-terminus, most notably M1, coverage in the remaining regions is near complete for all three insertion datasets (**Fig. 2**).

**Figure 2:**
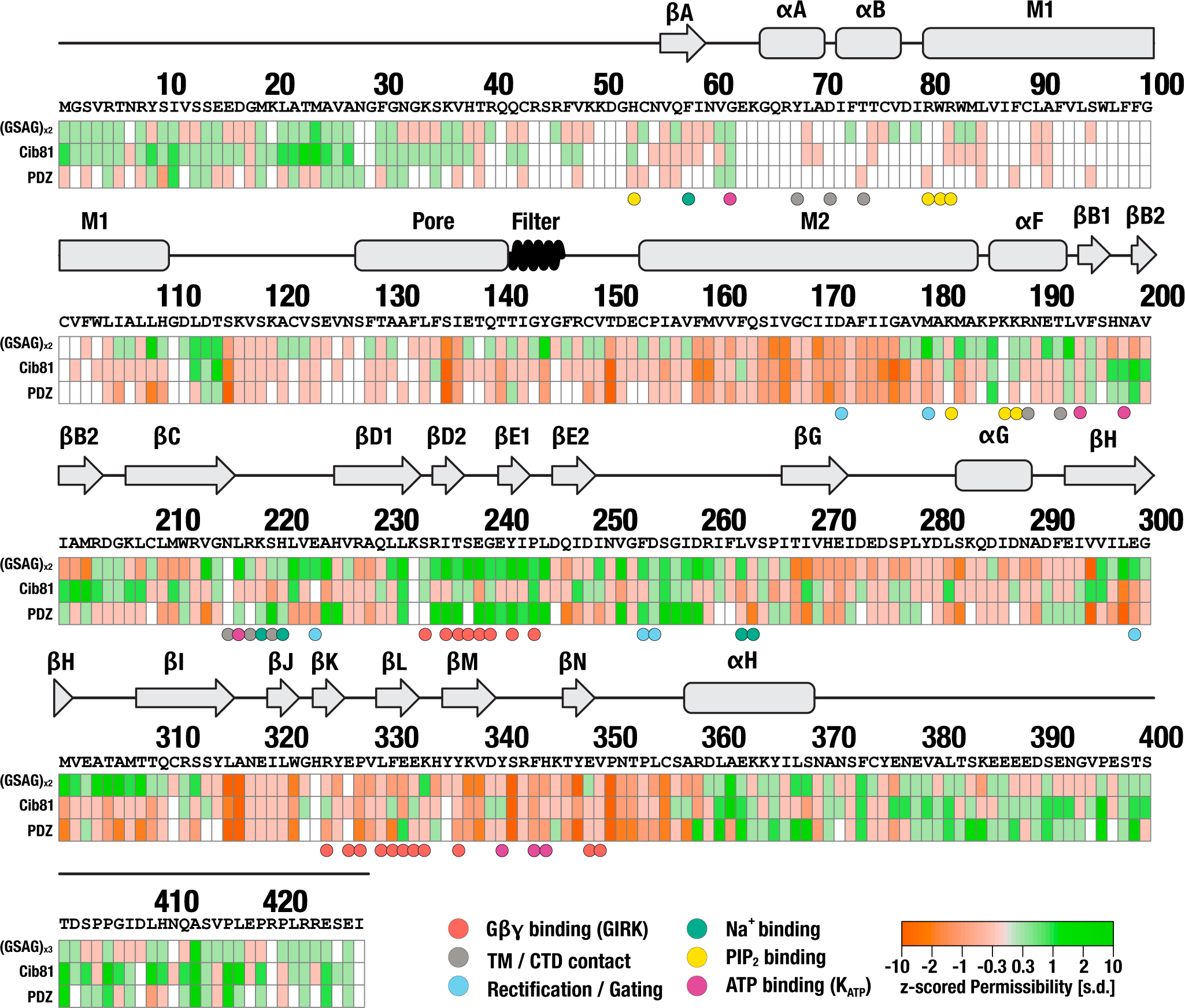
Domain Insertion Permissibility in hKir2.1. The primary sequence of human Kir2.1 (GI: 4504835) and secondary-structure elements are shown along the permissibility score for three types of inserted domains (indicated on the left). Key residues with functional relevance are indicated by color-coded spheres below.

### Domain insertion permissibility is surprisingly different between domains

We then mapped permissibility onto the crystal structure of chicken Kir2.2 (PDB 3SPI)^34^. As expected, domain insertion positions that should not allow surface expression do not (e.g., transmembrane and inter-subunit interfaces, **Fig. 3a, Supp. Fig. 3–5)**. Unsurprisingly, the unstructured C-terminus (which *in vivo* interacts with scaffolding proteins not present in HEK293 cells) was highly permissive to any insertion (**Fig. 2**). Predictably, overall flexible peptide insertions are more permissive than larger, more structured domains. Counter-intuitively, many non-conserved, surface-exposed loops (e.g., αG-βH or βN-αH) were also not permissive (**Fig. 2, Fig. 3b**). From this, we conclude that the rules that govern permissibility, at least in Kir2.1, differ from cytosolic proteins. Perhaps this is due to more stringent selection pressures unique to membrane proteins such as the need for proper folding, assembly, surface trafficking, and membrane insertion.

**Figure 3:**
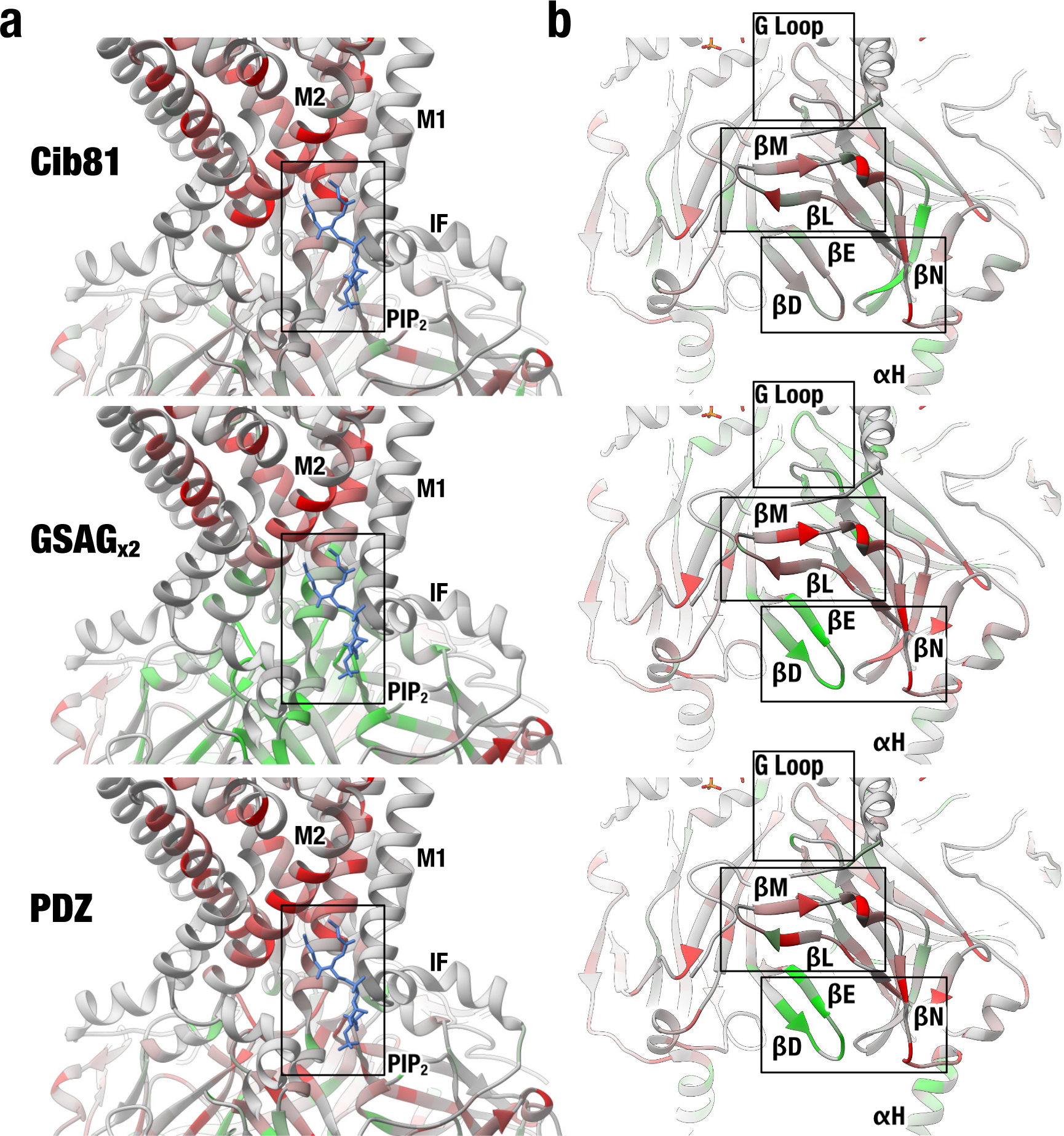
Differential Domain Insertion Permissibility. Permissibility data is mapped on the crystal structure of chicken Kir2.2 (PDB 3SPI). Insertion permissibility for three different domains (indicated on the left) is shown. **(a)** All types of insertions into transmembrane helixes and intersubunit interfaces are strongly selected against. Permissibility in PIP2 binding site depends on structural context of the insertion. (**b**) many non-conserved, surface-exposed loops (e.g., βG-αH or βM-βN) were not permissive, while the βD-βE loop (which binds Gβγ in GIRK) and the G loop (the cytoplasmic gate in Kir2.1) have context-dependent permissibility.

We more quantitatively compared permissibility profiles of biological replicates of Kir2.1 for inserted domains (structured vs. flexible) by clustering correlation matrices (**Supp. Fig. 6a**). Biological replicates were in very good agreement, which indicates that relatively little noise was introduced through the transposition, heterologous expression, and cell sorting steps. We found that domains cluster by structure (Cib81 and PDZ cluster discretely from each other, but flexible insertions do not) and size (Cib81 and PDZ cluster closer than flexible peptides).

### Allosteric Sites are most differentially permissive between inserted domains

Our data also reveals that permissibility is sensitive to the structural context of the inserted domain. Despite Cib81 and PDZ being of similar size, many sites are *differentially* permissive, suggesting that there is context dependence for permissibility beyond simple sterics (**Fig. 2 & 3**). Further demonstrating permissibility’s context dependence, while overall flexible linkers had highest permissibly there are several sites where only Cib81 and PDZ are tolerated (βB2-βC). Surprisingly, all of Kir2.1’s known allosteric sites and many regulated sites are differentially permissive. This included the PIP_2_ binding site at the interface between the pore and cytosolic domain (**Fig. 3a**), and the βG loop, a flexible region involved in channel opening (**Fig. 3a**). Particularly interesting is that the remaining highly yet differentially permissive sites –outside unstructured termini– are only known to be allosterically regulated in homologs of Kir2.1; they are thus putative latent allosteric sites. This includes the βB-βC loop (which binds ATP in Kir.6x ^36^) (**Fig. 2**), and the βD-βE loop (which binds Gβγ in GIRK^43^) (**Fig. 3a**). Correlation matrices of permissibility profiles sorted by insertion sites reveal distinct clusters that coincide with structural features involved in allosteric regulation of Kir2.1 (**Supp. Fig. 6b, 7**). Permissibility in exploited allosteric sites in Kir2.1 (the PIP_2_ binding site and G loop) is more strongly correlated than in putative latent allosteric sites (βB-βC, βD-βE, and βL-βM loops).

### Domain insertion permissibility is primarily dependent on dynamic protein properties

Given the unexpected patterns of domain insertion permissibility, we explored what protein features most contribute to permissibility. We calculated and compared structure-, conservation, and dynamic-based properties for Kir2.1 with different domain permissibility profiles (**Fig. 4a**). We found that domain insertion permissibility is correlated with dynamic features and does not correlate well with static and conservation-related protein properties.

**Figure 4:**
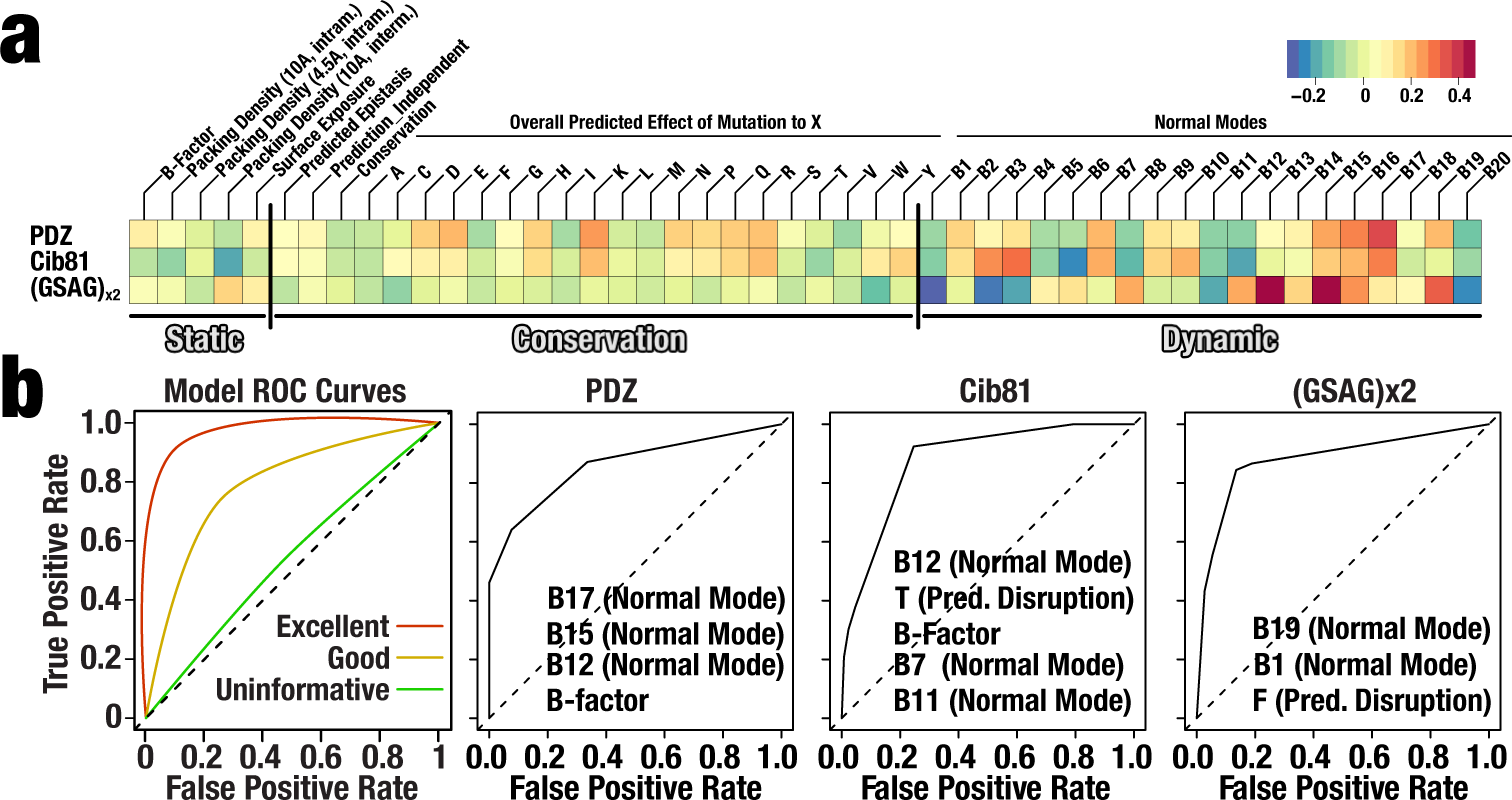
Parameter Correlation and Model Performance. **(a)** Spearman correlations between permissibility for the inserted domain indicated left with the calculate property indicated on the top. Vertical bars separate feature category (Static, Conservation, and Dynamic). **(b)** Model performance determined with receiver operator characteristic (ROC) curves for the indicated domain, along with the top predictive properties used in each recursive 10-fold cross-validated decision tree model. Examples of ROC curves for models with different performance are shown on the left-most panel.

To determine the degree to which computed properties can explain permissibility, and which properties best approximate permissibility, we constructed decision tree classification models. Consistent with the result that permissibility best correlates with dynamic properties, dynamic features had the greatest predictive power across the three types of domains (**Fig. 4a, Supp. Fig. 8-10**). While some profiles’ predictive models perform better than others (GSAG_x2_ was best and PDZ was worse) and none fully explain permissibility, we *can* build models for all domains, whose performance is far better than random as assessed by receiver operating characteristic (ROC) curves and other performance criteria. Our ability to generate predictive models for permissibility demonstrates that features used in predicting measured permissibility are meaningful and that the ‘indefinable qualities’ of permissibility play minor roles. Furthermore, the necessity for a non-linear approach (decision trees) demonstrates that permissibility, and by extension latent allostery, is an emergent phenotype from non-linear interactions between multiple protein properties, as opposed to a linear combination of, for example, conservation and surface exposure.

### Domain insertions in allosteric sites also have context dependent impact on function

Since permissibility only reports on whether a Kir2.1 insertion variant can fold and traffic to the cell surface, we focused on a representative sample of insertion positions to assess whether they remain functional (i.e., able to conduct K^+^) upon domain insertion. We subjected this subset to a flow cytometry-based optical activity assay that measures population-level resting membrane potential (RMP) in HEK293FT cells using an oxonol voltage-sensor, DIBAC4(3) (**Fig. 5a**)^44^. Since Kir2.1 drives the RMP towards the reversal potential of K^+^, cells expressing functional Kir2.1 are more hyperpolarized compared to ‘empty’ cells under our culture conditions (**Fig. 5b**). As expected, insertions into flexible and highly permissive regions (e.g., M24, A371) of Kir2.1 have little impact on function. All insertions into regions critical for gating, the G-loop or the conduction pore exit, break channel function. Whereas, insertions into the exposed extracellular loop (S116) have different impacts on function. Here, both Cib81 and a flexible linker are well tolerated, while PDZ impairs function significantly. The emerging theme of differential impact on function continues in other regions of Kir2.1, including the PIP_2_ binding site. Here, P187 is permissive to all insertions, but partial function remains with large domain insertions (PDZ & Cib81) while a flexible linker completely breaks channel function. Conversely, permissibility and impact on function tracked quite well for insertions into the D-E loop. Here, both PDZ and flexible insertions allowed gating, while Cib81 broke channel function. Overall as with permissibility, the functional assay shows that domain insertions in allosteric sites impact function in a context dependent-manner that cannot be explained with simple sterics.

**Figure 5:**
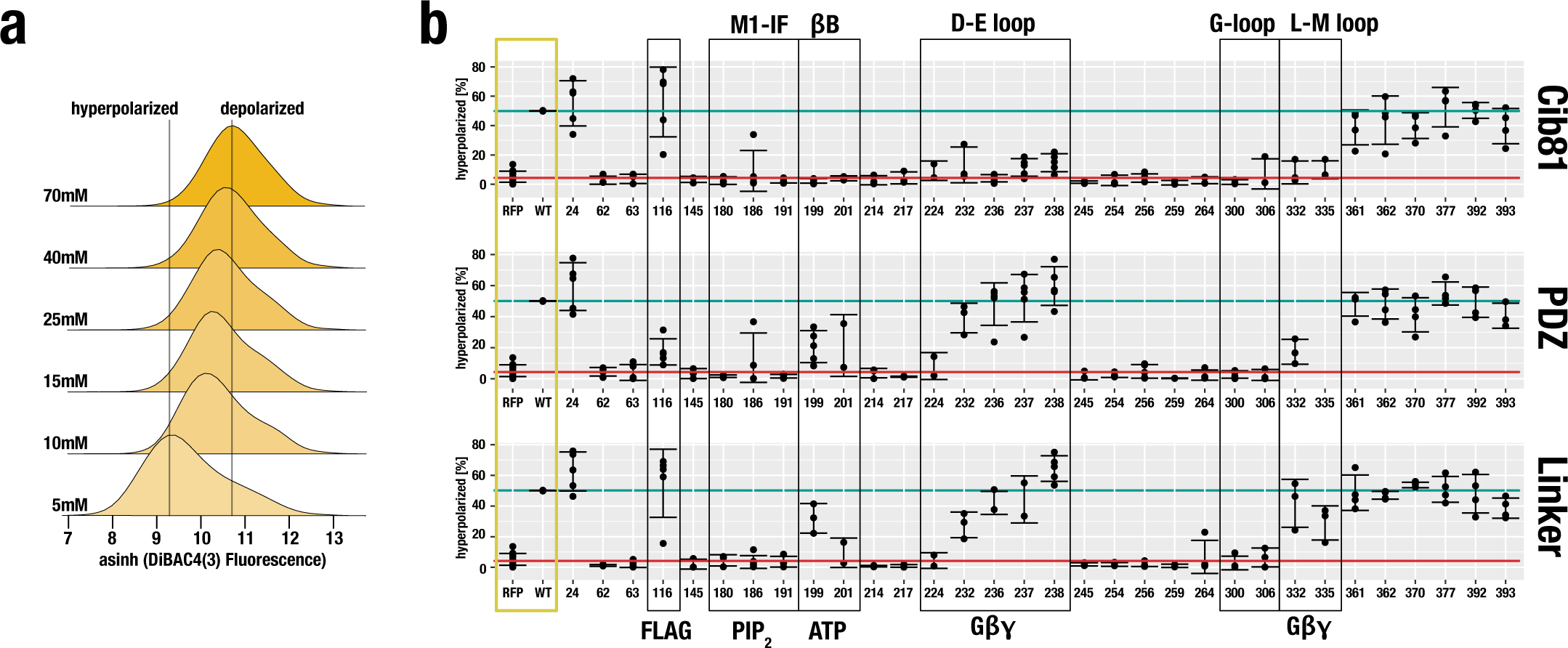
Domain Insertion Impact on Kir2.1 Function. **(a)** Population-level DiBAC4(3) fluorescence in HEK293FT cells expressing Kir2.1 as a function of external K^+^. Increasing K^+^ depolarizes the cells, resulting in less membrane-partitioning of the dye thus increasing measured fluorescence. **(b)** Shown are the percent hyperpolarized cells expressing the indicated Kir2.1 variant and inserted domain. Higher numbers indicate function, and lower numbers indicate disruption. Reference measurements are provided for HEK239FT expressing miRFP670 alone and wildtype Kir2.1 co-expressed with miRFP670 (yellow box). Reference levels of WT and no channel are indicated by horizontal green and red lines, respectively. Regions discussed in the text are indicated by black boxes.

### Switchable domain insertions in allosteric sites modulate channel function

What permissibility and functional assays tell us is that many allosterically regulated sites are sensitive to the structure of the inserted domain (differentially permissive) and retain at least partial function. The fact that sites involved in allosteric signaling in Kir2.1 homologs (GIRK and Kir6.2) have similar features suggests that these sites might have latent allosteric potential. To test this idea, we tested insertion Cib81 variants in these sites for light-dependent modulation, as well as several others that we predicted are non-allosteric (e.g., C-termini), a sterically inaccessible extracellular loop, and negative controls (wildtype and poor gating mutant, V302M ^45^). Initially, there was no optimization of flanking linkers. We reasoned that if Cib81 is sterically accessible and a position has allosteric potential, then a light-mediated association of the channel with co-expressed Cry2 (size 70 kDa) would modulate channel gating even if binding interfaces are not optimized. We adopted the flow cytometry RMP assay to measure Kir2.1 activity with and without blue light illumination. As expected, wildtype channel and a gating mutant have no light-dependent modulation. Furthermore, when Cib81 is inserted into predicted non-allosteric sites terminal regions (e.g., E392) recruitment of Cry2 to the channel does not affect Kir2.1 activity. Similarly, when Cib81 is inserted into an inaccessible extracellular site (S116), there was no light-dependent effect on Kir2.1 activity. Remarkably, even though channel function was severely impaired, when Cib81 was inserted in the PIP_2_ binding site, illumination markedly decreased the remaining Kir2.1 activity (**Fig. 6a-b**). We validated this with patch clamp electrophysiology, which shows that the open probability of Kir2.1(P187CIB), which is low to begin with, is further *decreased* with blue light illumination (**Fig. 7a-b**).

**Figure 6:**
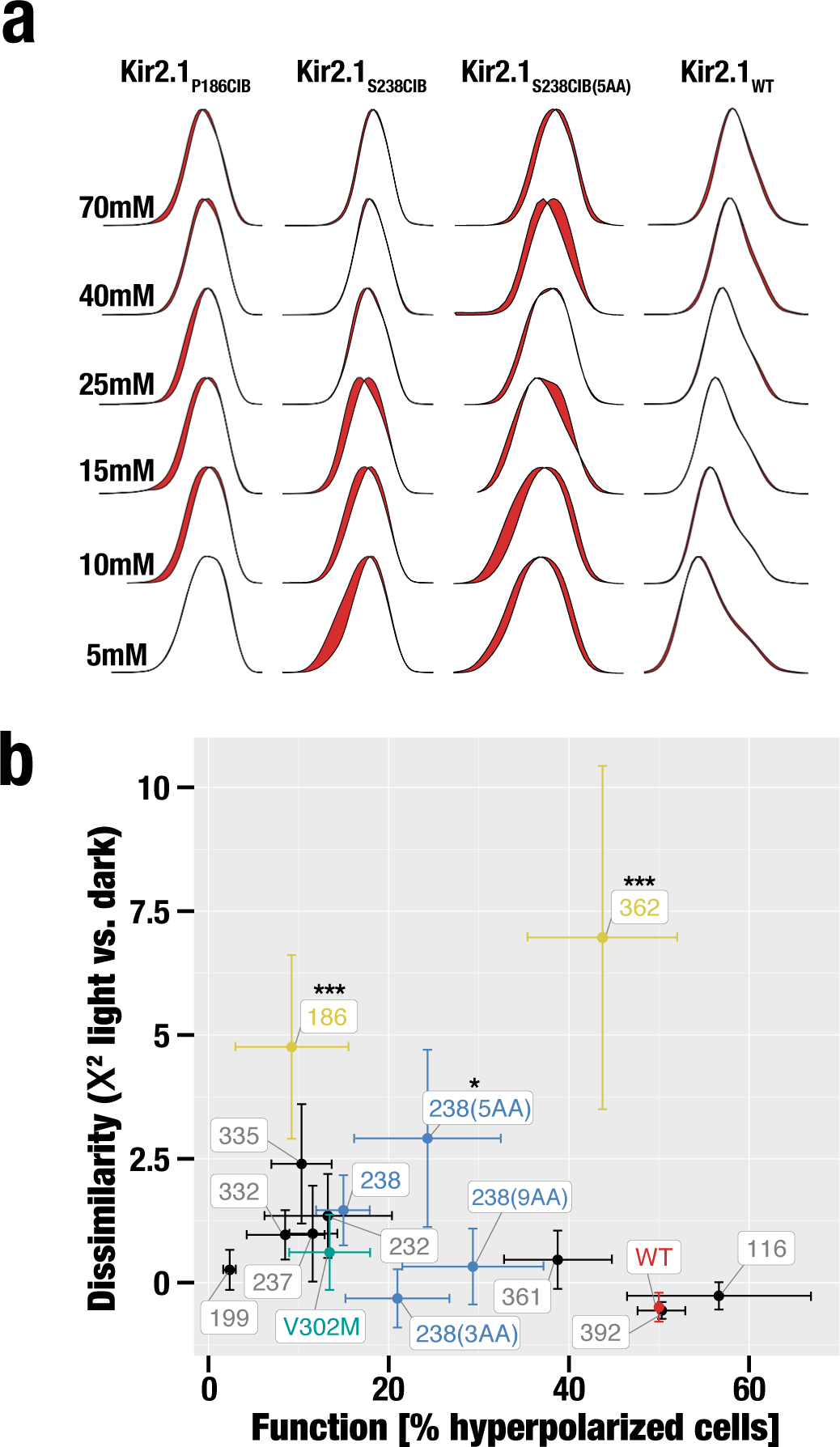
Light-modulated Kir2.1 Variants. **(a)** Representative examples showing the difference (indicated by red areas) in population-level DiBAC4(3) fluorescence with and without blue light illumination for HEK293FT cells expressing the indicated Cib81 insertion mutant or WT channel and Cry2. **(b)** Quantitation of dissimilarity (X^2^) compared to channel function. Highlighted are insertions into the PIP_2_ binding site and αH helix (yellow), D-E loop (blue), gating mutant V302M (green), and wildtype Kir2.1 (red). Significance of light modulation is tested by pairwise comparisons using Dunnett's test for multiple comparisons with wild type as control and post-hoc multiple comparison adjustment. Significance levels: *** p < 0.001; ** p < 0.01, p > 0.05 (not significant)

**Figure 7:**
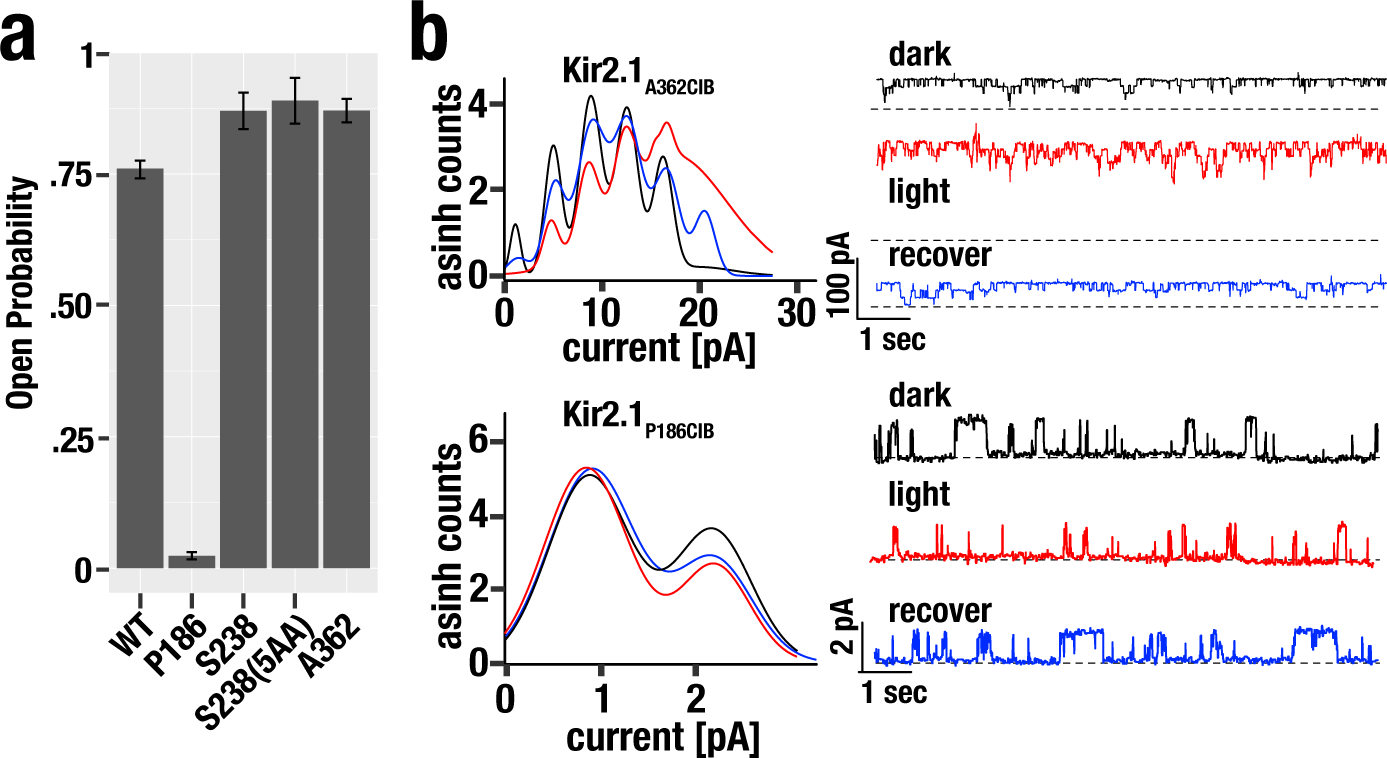
Electrophysiology of Light-switched Kir2.1 Variants. **(a)** Open Probability determined by on-cell patch clamp electrophysiology for *wt* Kir2.1 or the indicated insertion mutant. Error bars are s.e.m. (n = 3–12). **(b)** Gaussian functions (Kir2.1_A362CIB_, k = 6; Kir2.1_P186CIB_ k = 2) fit to all points current amplitude histograms for representative examples of the indicated insertion mutant. Before, during, and recovery (>2 min post-illumination) are indicated by black, red, and blue lines, respectively. Representative current traces are shown on the right. Dashed lines indicate zero current level.

We also observed Kir2.1 light-modulation when Cib81 was inserted into the pore-facing side of the αH Helix (A362), but not the outward-facing side (L361) (**Fig. 6b**). Furthermore, patch clamp validation of this insertion showed that open probability is higher than wild-type channel in the absence of illumination (**Fig. 7a**) and further *increased* with illumination (**Fig. 7b**). The αH Helix is a potential Gα binding site in GIRK based on several NMR structures; however, this interaction hasn’t been fully explored^46^.

Most interestingly, we noticed weak light modulation when Cib81 was inserted into predicted latent allosteric sites which are part of the βD-βE and βL-βM loops (S238, L332, R335) (**Fig. 6b**). The weak impact of Cry2 recruitment in these regions could be due to none-optimized binding interfaces in contrast to Gβγ to GIRK’s βD-βE and βL-βM loops^43^. When we patched cell expressing one of these insertion mutants (S238), we observed higher Kir2.1 activity again even in the absence of illumination (**Fig. 7a**), suggesting that insertions into the βD-βE loop are activating. We explored linker optimization to obtain a cleaner photoswitching phenotype. Flanking the inserted Cib81 by five amino acids, but not three or nine, improved light modulation of Kir2.1 significantly (**Fig. 6b**).

## Discussion

Our findings reveal that permissibility in ion channels is correlated with protein dynamics, but not structural features or conservation metrics. There is broad support for the idea that protein dynamics provides a mechanism through which a new function can come under natural selection^21,22^. At the same time, for this conformational flexibility to be exploited for allosteric regulation, it must be constrained and not too degenerate; that is, it must be sensitive to the context of the perturbation caused by an allosteric modulator for there to be specificity. Translated into the language of permissibility this means we expect that ‘unimportant sites’, e.g., those found in disordered terminal extension to be permissive of any insertion without affecting the functional phenotype of the channels (**Fig. 8a**). Structurally important sites, e.g., those forming the pore of an ion channel, are expected to cause the misfolding and a loss of function phenotype regardless of the inserted domain’s properties. Sites that have conformational plasticity that depends on the context of a perturbation, e.g., sites involved in either exploited or latent allostery, will have *differential* permissibility. This means that the effect of a domain insertion on both folding, assembly, and trafficking will depend on the biophysical properties of the inserted domain. In a similar vein, depending on the context of the inserted domain, different functional phenotypes are expected. We have found sites belonging to each of these three categories in Kir.2.1.

**Figure 8:**
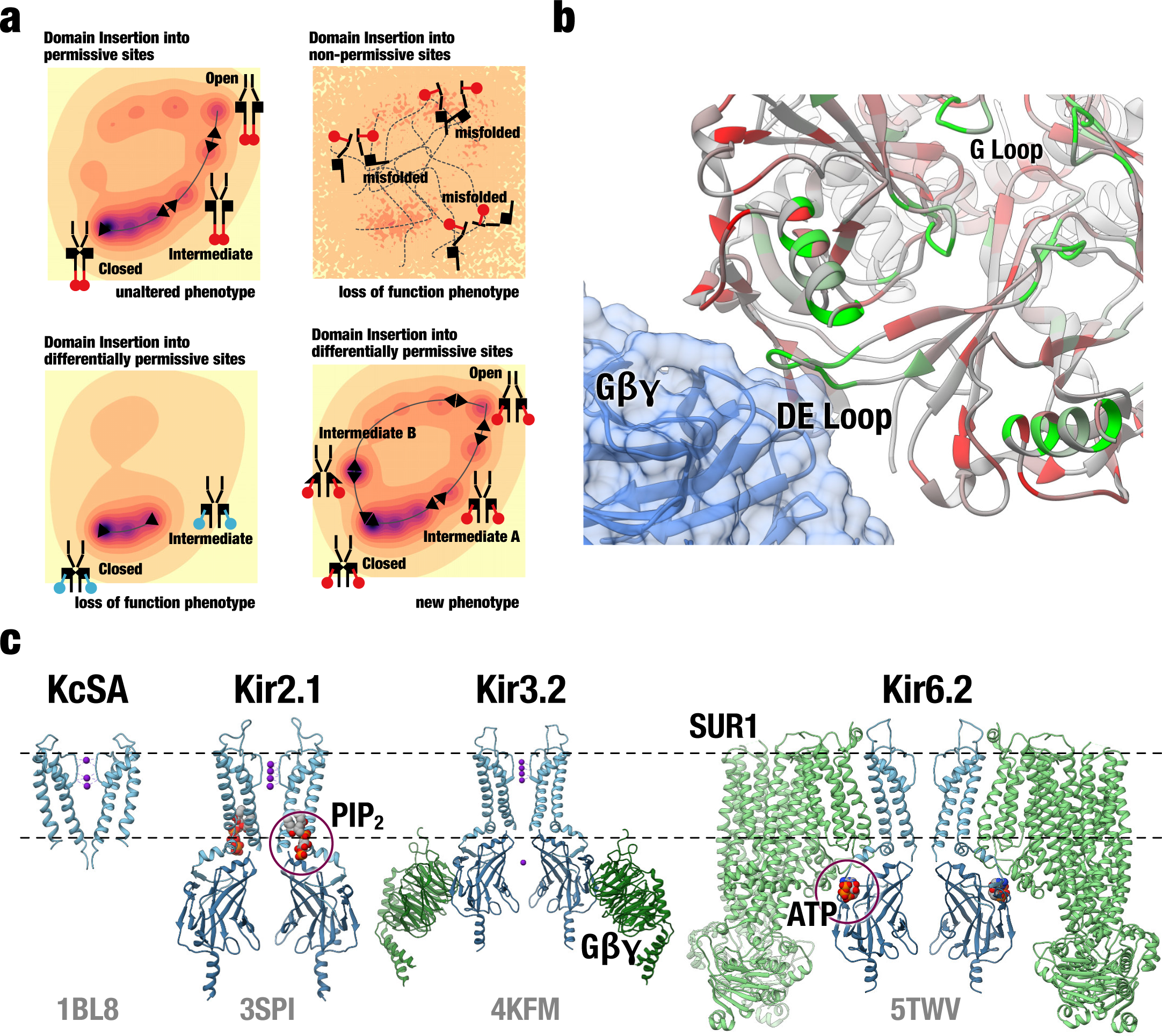
The Role of Conserved Channel Architecture in the Evolution of Divergent Allosteric Regulation. **a)** Cartoon model that illustrates the different outcomes of domain insertion. Insertions into highly permissive sites will have little impact on folding and function. Insertions into structurally important sites (non-permissive) will interfere with folding and cause a loss of function. Insertions into sites involved in conformational transition required for allosteric regulation will have context dependent permissibility and are associated with loss of function or gain of a new function. **(b)** Domain Insertion Permissibility mapped onto the crystal structure of GIRK2 (Kir3.2) in complex Gβγ (PDB 4KFM). Many highly permissive sites in Kir2.1 are homologous to those that interact with Gβγ in GIRK2. **(c)** Domain structure and structural conservation between K^+^ channels. Pore domains are shown in light blue, intracellular domains are shown in dark blue. Small molecule allosteric modulators (if applicable) are shown as space-filling models. Proteinous allosteric modulators are shown in light and dark green. Remarkably, the same overall architecture is exploited for different allosteric modulation modes.

Examples of the first category, ‘unimportant sites’, are many sites toward the C-terminus, which have *high* permissibility to any domain insertion and where the impact on function is minimal, irrespective of insertion type. Universal permissibility likely means that these regions play minor roles in surface trafficking, oligomerization, or channel activation. This is consistent with their known role for interacting with binding partners, most of which are not present in HEK293 cells^47-^49. Interestingly, domain insertions into the F_374_CYENE motif for Golgi export that is present in the C-terminus of Kir2.1 were reasonably well tolerated. Previously, it was reported that even conservative mutations in this motif essentially abolish surface expression of GFP-tagged Kir2.1^50^. One possible explanation is that a domain *insertion* does not replace this motif; it only becomes discontinuous at the primary sequence level, while retaining tertiary structure. Another explanation is that the F_374_CYENE motif has 'dominant phenotype' in the sense that only one monomer providing an unperturbed copy is sufficient for the channel to be exported and surface-trafficked. In our assay– insertion libraries transfected into HEK293 cells– channels can assemble from monomers with different insertions, and we have not explicitly investigated 'copy number' effects of domain insertion permissibility. We do not yet know if either effect explains our experimental data; the larger point raised by this example is that we have only begun to understand what insights we can gain from these types of experiments. Further studies, in particular with more diverse sets of inserted domains and including purifying rounds of selection, are needed to establish a more fine-grained picture of the impact of domain insertions.

Examples for the second category, scaffold sites, include transmembrane helices, which have *low* permissibility to any insertion, and where any domain insertion severely impacts function. The universal disruptiveness of mutations within these regions is likely due to these regions being essential for folding, oligomerization, surface trafficking or membrane insertions. This is consistent with many scaffold sites occurring within transmembrane domains or at interfaces between channel monomers. One of the sites in this category that was among the least permissive to domain insertions is the S_314_YLANEILW Golgi export motif in the βI-βJ loop. Mutations in this motif, in particular, Kir2.1Δ314-315, are associated with the Andersen-Tawil Syndrome^51^. The severity of any domain insertion into this motif is unlike what we observed for the F_374_CYENE motif. Perhaps this can be explained by this region forming a specific tertiary structure that must interact with the RXR_46_XXK motif in the N-terminus to prevent accumulation in the Golgi^52^. This potential explanation is strengthened by the fact that the S_314_YLANEILW motif is within a highly-structured region whereas F_374_CYENE is in an unstructured region of Kir2.1.

Sites in the third category have *differential* permissibility that depends on both the structural context of the insertion position as well as the biophysical properties of the inserted domain. In these sites, we find that impact on function is not correlated with permissibility. We postulate that these sites are exploited or latent allostery sites. Indeed, in addition to the binding pocket of Kir2.1’s allosteric modulator PIP_2_, significant differential permissibility was measured in the βD-βE and βL-βM loops. While these loops have no known regulatory function in Kir2.1, they are part of an alcohol-binding pocket conserved in both Kir2.1 and GIRK^53,54^. Functional analysis in GIRK revealed that alcohol modulates PIP_2_-mediated channel activation in a G-protein independent way ^55^. Furthermore, the βD-βE and βL-βM loops are critical for mediating activating interactions between Gβγ and GIRK (**Fig. 8b**)^43^. That inserting PDZ or Cib81 domains, whose potential interaction surface is roughly similar to that of Gβγ’s, into this loop resulted in an activating phenotype suggests that Gβγ modulation of GIRK can perhaps be thought of as two different processes: One is specific binding mediated by residues that exist in GIRK but not in Kir2.1, and the second being a mechanism for coupling this binding to channel opening that exists both in GIRK to a lesser extent in Kir2.1 because of the shared architecture of the C-terminal domain (CTD). This type of division of labor, where one set of sites encodes affinity, while the other set encodes a filter for efficacy has been described for several types of allosteric regulation, including Gα activation of GPCRs^56^ and ligand binding to bioamine receptors^57^.

It is well-established that allostery and latent phenotypes emerge primarily from dynamic protein features ^21,22^. That permissibility is best correlated and predicted by dynamic properties and not those describing static structural and conservation features of amino acids, is therefore entirely in agreement with the proposed mechanism through which dynamic protein properties enable latent allosteric properties to be exploited in response to novel regulatory factors^24^. If a protein can dynamically switch between ground and excited states, then the protein can retain its original ground-state native structure, while at the same time stabilizing an excited state structure incrementally, thus providing continuous viability during evolutionary transitions^58^. Our ability to build predictive models of permissibility also means that if models are trained on a sufficiently large experimental dataset, it might be possible to derive generalized predictive models that can predict permissibility on potentially any ion channel, thus rendering case-by-case mapping of permissibility superfluous. It will be interesting to see if permissibility can be predicted from the same set of calculated properties for any ion channel (indicating permissibility is universal), or whether it is a function of phylogenetic distance (indicating permissibility is ‘path dependent’).

Interpreted broadly, mapping and building models of permissibility– and by extension allostery–as it changes through phylogeny may be useful in explaining how specific ion channel families evolved. Much of the core functionality and architecture of ion channel families had evolved by the time the metazoan lineage appeared^29,30^. Subsequent diversification, driven by adaptive pressure to develop specialized neuromuscular tissue, can be considered fine-tuning biophysical properties and evolving novel regulation. We can observe this in K^+^ channels (**Fig. 8C**). After the inward rectifier architecture (represented by Kir2.1) evolved from the simpler pore-only architecture (KcSA) by the addition of the CTD, the same overall architecture is utilized for different modes of allosteric regulation, including different small molecule ligands (PIP_2_, ATP, Na^+^) and proteins (Gβγ and SUR1). The notion of latent allostery can explain how this came to be. Because the dynamic features for the most part arise from global architecture fine-tuned by local interactions^59,60^, and because potential for allostery is an emergent by-product of these dynamics features^22^, it is entirely expected that a new allosteric regulation scheme leverages a pre-existing pathway and can evolve without gross changes to overall topology.

To more practical interest, we demonstrate that inserting switchable domains only into existing allosteric sites or predicted latent allosteric sites, renders the target channel light-switchable. We do not yet know whether this is generalizable to multiple ion channels. However, what this proof-of-principle experiment demonstrates is that mapping latent allosteric sites via experimentally measured permissibility could greatly simplify engineering ion channels with useful function, such as sensitivity to biorthogonal stimuli.

## ACKNOWLEDGEMENTS

We thank Matthew R. Whorton, Mikael Elias, Sivaraj Sivaramakrishnan, and the entire Schmidt lab for helpful feedback and discussion, Therese Martin with flow cytometry technical advice and support, David Savage and Avi Flamholz for technical advice and development of the DIPseq alignment pipeline, Alina Zdechlik, Tejas Gupte, and Daniel Sorenson for helpful feedback on the manuscript, and Steffan Okorafor for assistance with single mutant construction. W.C.M. is funded by a National Science Foundation Graduate Research Fellowship.

## Author Contributions

W.C.M. and D.S. conceived the study. W.C.M. conducted domain insertion permissibility experiments, predictive model building, flow cytometry functional assays, and flow cytometry optogenetic assays. Y.H and D.S. conducted electrophysiology experiments. C.L.M provided expertise for predictive model building. W.C.M. and D.S. co-wrote the manuscript.

## Supplementary Materials for

### Materials & Methods

#### MuA domain insertion library generation

Transposition libraries were generated using 100ng MuA-BsaI engineered transposon and 1:2 molar ratio transposition target DNA in 20ul reactions with 4ul 5x MuA reaction buffer and 1ul 0.22 ug/ul MuA transposon (Thermo Fisher). MuA-BsaI engineered transposon propagation plasmid or pUCKanR-Mu-BsaI was a gift from David Savage (Addgene plasmid # 79769)^61^. MuA-BsaI engineered transposon was digested with BglII and HindIII Fastdigest enzymes (Thermo Fisher) and gel purified using gel purification kit (Zymo Research).

The transposition target, human Kir2.1 (GI: 4504835) including a porcine teschovirus ribosomal skipping sequence (P2A)^62^, was codon-optimized for mouse, synthesized (Gen9) and subcloned into pATT-Dest using NEB BamHI and HindIII. pATT-Dest was a gift from David Savage (Addgene plasmid # 79770)^61^. A FLAG tag was inserted after T115 using Q5 site-directed mutagenesis. MuA transposition reactions were incubated at 30degC for 18 hours for transposition, followed by 75degC for 10 minutes for heat inhibition. DNA from reactions was cleaned up (Zymo Research) and eluted in 10ul water. All 10ul were transformed into 30ul electrocompetent 10G ELITE E. coli (Lucigen) in 1.0 mm Biorad cuvettes using a Bio-Rad Gene Pulser II electroporator (settings: 10uF, 600 Ohms, 1.8 kV). Cells were rescued and grown without antibiotics for 1 hour at 37degC. Aliquots were then serially diluted and plated on LB agar plates containing carbenicillin (100 ug/ml) and chloramphenicol (25 ug/ml) to assess library coverage. The remaining transformation mix was grown in 50 ml LB containing carbenicillin (100 ug/ml) and chloramphenicol (25 ug/ml). All transformed libraries yielded greater than 10^5 colonies, which for Kir2.1-P2A (1369bp) is >35x coverage. Plasmid DNA was purified by midi-prep kit (Zymo Research).

Transposition-inserted Kir2.1 variants were subcloned into an expression vector by amplifying channel variant genes, adding on BsmbI sites, using 10 cycles of PCR using Primestar GXL (Takara Clontech) and run on a 1% agarose gel. The larger band was cut out and gel purified (Zymo Research) to isolate channels with inserted transposons. A mammalian expression vector (pcDNA3.1) with EGFP was amplified to add on BsmbI sites complementary to those on Kir2.1-P2A. The Kir2.1-P2A (BsaI-transposon) variants where subcloned into this vector by BsmbI-mediated Golden Gate cloning^63^. Reactions were cleaned (Zymo Research) and eluted with 10ul water. All 10ul were transformed into 30 ul Lucigen electrocompetent 10G ELITE E. coli and electroporated in 1.0 mm Biorad cuvettes using a Bio-Rad Gene Pulser II electroporator (settings: 10 uF, 600 Ohms, 1.8 kV). Cells were rescued and grown without antibiotics for 1 hour at 37degC then with an aliquot serially diluted plated on LB agar plates containing kanamycin (50 ug/ml) and chloramphenicol (25 ug/ml) to assess library coverage. The remaining transformation mix was grown in LB containing kanamycin (50 ug/ml) and chloramphenicol (25 ug/ml). All transformed libraries yielded greater than 10^5 colonies so for Kir2.1 (1369bp) there is >35x coverage. Plasmid DNA was purified by midi-prep kit (Zymo Research).

Inserted Transposons were replaced with domains in individual reactions using BsaI-mediated Golden Gate cloning^63^. Domains (PDZ, Cib81, GSAG_x2_, GSAG_x3_) for insertions were ordered as gblocks (IDT DNA), and PCR amplified to add on BsaI sites complementary to MuA-BsaI transposon sites for Golden Gate cloning. Domain amplicons were gel purified (Zymo Research). The product was further digested with AgeI-HF (NEB) and Plasmid-Safe ATP-dependent DNase (Epicentre) to remove any undigested transposon, then cleaned up (Zymo Research) and eluted with 10ul water. All 10ul were transformed into 30 ul Lucigen electrocompetent 10G ELITE E. coli and electroporated in 1.0 mm Biorad cuvettes using a Bio-Rad Gene Pulser II electroporator (settings: 10 uF, 600 Ohms, 1.8 kV). Cells were rescued and grown without antibiotics for 1 hour at 37degC. An aliquot was serially diluted and plated LB agar plates containing kanamycin (50 ug/ml) to assess library coverage. The remaining transformation mix was grown in LB containing kanamycin (50 ug/ml). All transformed libraries yielded greater than 10^5 colonies so for Kir2.1 (1369bp) there is >35x coverage. Plasmid DNA was purified by midi-prep kit (Zymo Research).

#### Domain insertion permissibility cell sorting assay

100 ng of each domain insertion library was transfected with 36 ul of turbofect (Thermo Fisher) into 50% confluent HEK293FT(Invitrogen) with 11.9 ug of dummy plasmid (pATT Dest) divided across a single 6 well dish (9.6 cm^2^ / well).

Cells from each well were detached using 1 ml accutase (Stemcell Technologies) and twice spun down at 450xg and resuspended in FACS buffer (2% of FBS, 0.1% NaN3, 1xPBS). Cells were incubated with 1:200 anti-flag mouse antibody (Sigma) 1 hour rocking at 4degC, washed twice with FACS buffer, covered with aluminum foil, and then incubated with 1:400 anti-mouse Alexa Fluorophore 568 (Thermo Fisher) for 30 minutes rocking at 4degC. Cells were washed twice, resuspended in 3 ml FACS buffers, and filtered using cell strainer 5 ml tubes (Falcon). Cells were kept on ice and protected from light in the transfer to the flow cytometry core. Before cell sorting, a small aliquot of cells was saved as a control sample for sequencing.

Cells were sorted into EGFP high / Alexa568 low (transfected cells without surface expression) and EGFP high / Alexa Fluorophore 568 high (transfected cells with surface expression) on a BD FACSAria II P69500132 flow cytometer. EGFP fluorescence was excited using a 488 nm laser, recorded with a 525/50 bandpass filter and a 505 long pass filter. Alexa fluorophore 568 fluorescence was excited using a 561 nm laser and recorded with a 610/20 bandpass filter. Cells were gated on Side Scattering and Forward Scattering to separate out whole HEK293FT cells, gated on forward scattering area and width to separate single cells, then gated on co-expressed EGFP to gate out cells that received a plasmid, then gated on cells that were labeled using the anti-flag antibody for surface expressed channels. Gates were determined using single wildtype, EGFP only and unstained library samples. EGFP high / Label low and EGFP high / Label high cells were collected into catch buffer (20% of FBS, 0.1% NaN3, 1xPBS). Between 2,000-100,000 cells were collected for each sample/library pair which is ~4-250x coverage of all potentially productive (i.e., in-frame and forward) domain insertions.

DNA from Control, EGFP high / Label low, and EGFP high / Label high cells for each library were extracted using a Microprep DNA kit (Zymo Research) and triple eluted with water. To remove chromosomal DNA, samples were digested with Plasmid-Safe ATP-dependent DNase (Epicentre). The resulting plasmid DNA was further purified and concentrated using (Zymo Research). The product was used as a template for 12 cycles of PCR using Primestar GXL (Takara Clontech), run on a 1% agarose gel, and gel purified (Zymo Research) to remove any primer dimers or none amplicon DNA. Purified DNA was quantified using Picogreen DNA concentration at the University of Minnesota Genomics Core. Equal amounts of each domain insertion sample were pooled by cell sorting category (control, EGFP high / Label low, EGFP high / Label high were pooled for sequencing library generation and sequencing.

#### Sequencing

Libraries were generated at University of Minnesota Genomics Core using Nextera XT or Nano Truseq library generation (Illumina) to fragment and add on Illumina sequencing adaptors and sequenced using either HISEQ or MISEQ sequencing platforms.

#### Domain insertion permissibility alignment and enrichment

Alignments were done on both forward and reverse reads using a DIPseq pipeline developed by David Savage and coworkers that we slightly modified for compatibility with updated python packages^40^. Reads with duplicate domain insertion calls in both forward and reverse reads were removed. This pipeline results in plaintext files indicating a domain insertion positions and whether that insertion is in-frame and in the forward direction. Enrichment was calculated by comparing the change in EGFP high /Label low to EGFP high / Label high cells. Only positions with reads in both samples were used in enrichment calculations. All these positions are treated as ‘NA’ and not considered in downstream analysis and structure mappings, with the exception of calculating correlations between datasets and correlations between sites. In these correlation calculations treat ‘NA’s as ‘0’s so removing all the data will introduce more noise when comparing between datasets due to limits from sampling.

Fitness function for individual datasets:

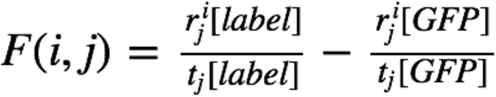

Where *r* is the number of reads at amino acid position *i*, in the *j*th dataset divided by *t*, the total number of reads in the *j*th given sample. This resulting data from individual sequencing reads are only used to calculate correlations between domains and amino acid positions.

For structure mappings and predictive model training means of fitness for a given domain insertion variant are used. So, the resulting mean fitness function is:

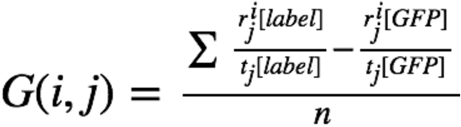

Where *r* is the number of reads at amino acid position *i*, in the *j*th dataset divided by *t,* the total number of reads in that given sample, summed for all replicates of that domain-channel combination, and divided by *n*, the number of datasets.

Differential fitness was calculated as the mean fitness standard deviation between Cib81, PDZ and GSAG_x1_ datasets.

Mean fitness and differential fitness was z-scored and mapped onto the structure of chicken Kir2.2 (PDB 3SPI) using Chimera ^64^. Mapped dataset for Cib81, PDZ and GSAGx1 linker had adequate coverage: 76.6% Cib81(333/435), 68.9% PDZ (300/435), and 76.7 % (334/435) of potential amino acid positions.

#### Dataset Correlations

Pearson correlations were used to calculate correlations between domain insertion datasets. Pearson correlations were also used to calculate correlations between amino acid positions across all datasets. Both these correlation matrices were calculated using the dataset that was trimmed to avoid sampling problems such that no more than 0.375 (6/16) datasets are where raw fitness was +10E-4 <x< -10E-4. These correlations were calculated with 63% of possible positions (277/435).

Spearman correlations were used to compare mean domain insertion datasets and calculated protein properties because Spearman correlations are often better at handling non-parametric correlations. Based on lack of structural and conservational data at various amino acid positions many sites had to be trimmed. Data was trimmed from positions where more than half datasets had a raw mean fitness was +10E-4 <x< -10E-4. This resulted in datasets that contained 70% of possible positions (207/293).

#### Computed protein properties

Static protein properties (B-factor, 10 Angstrom intramonomeric packing density, 4.5 Angstrom intramonomeric packing density, 4.5 Angstrom intermonomeric packing density, and surface exposure) were calculated using the SWIFT web server on chicken Kir2.2 (PDB: 3SPI)^34,65^, conservation based properties (overall predicted disruptive effect of a mutation, conservation, and individual predicted disruptive effect of a mutation to specific amino acids (A, C, D, E, F, G, H, I, K, L, M, P, Q, R, S, T, V, W, Y)) were acquired from the EVmutation data server^14^, and dynamic properties (first 20 normal modes of the B monomer) were calculated on the iGNM 2.0 normal mode webserver^66^. After trimming, computed protein properties were calculated for all parameters for 67% (293/435) of possible positions.

#### Decision tree models

We chose decision trees for building predictive models due to their utility in handling and determining non-linear interactions. Prior to training models, data was binarized such that 0 was not permissive and 1 permissive for a given domain insertion. After trimming any data for a given mean raw fitness between +10E-4 <x< -10E-4 datasets models were trained on 62.5% (183/293) Cib81, 58.0% (170/293) PDZ, and 68.6% (201/293) of possible positions. Models were limited to a depth of 4 to minimize model overfitting, trained on the computed protein properties to predict Cib81, PDZ, and GSAG_x2_ using the rpart package in R^67^, and cross-validated 10 times. Model performance was determined using commonly used criteria: receiver operating characteristic (ROC) curves, the complexity parameter (used in minimizing tree size)/tree depth and model residuals, precision vs. recall, accuracy vs. cutoff, precision vs. cutoff, and recall vs. cutoff (**Supp. Fig. 10**). As further validation, models were trained only the four most significant protein properties based on Spearman correlations to demonstrate necessity and utility of using decision trees vs correlation calculations (**Supp. Fig. 9A),** by withholding the most important properties determined with decision tree and by withholding whole classes of protein properties (static, conservation, dynamic, **Supp. Fig. 9B**).

#### Resting membrane potential functional assay

Single mutants were generated by inserting a BsaI site and a 5 basepair replication identical to those created by transposons that replicated the beginnings and ends of transposon-mediated domain insertions using a Q5 site-directed mutagenesis kit (NEB). Single insertion mutants were created for 32 sites (Amino Acid positions: 24, 62, 63, 116, 145, 180, 186, 191, 199, 201, 214, 217, 224, 232, 236, 237, 238, 245, 254, 256, 259, 264, 300, 306, 332, 335, 361, 362, 370, 377, 392, 393) and then replaced with the domains for which libraries were previously generated. Subsequently, using BsrGI and PstI sites, EGFP was replaced with miRFP670 for all mutants. miRFP670 was amplified from pmiRFP670-N1, which was a gift from Vladislav Verkhusha (Addgene plasmid # 79987)^68^. The same cloning approach was used to add 3-9 amino acid GSG linkers on either side of Cib81 in the 238 position.

The resting membrane potential assay was conducted on all aforementioned domain insertion mutants in addition to miRFP670 alone (negative control) and wildtype Kir2.1 (positive control). 400 ng of each mutant was transfected with 6 ul Polyethyleneimine (Polysciences) along with 600 ng of dummy plasmid (pATT-Dest) across 2 wells of a 24 well dish. For each experiment, wildtype Kir2.1 was transfected as a benchmark and for comparison for mutant function. Cells from each well were detached using 300 ul accutase (Stemcell Technologies), spun down at 450xg three times, and re-suspended in 200 ul Tyrode (125mM NaCl, 2mM KCl, 3mM CaCl2, 1mM MgCl, 10mM HEPES, 30mM glucose, pH 7.3). Bis-[1,3-dibutylbarbituric acid] trimethine oxonol (DiBAC4(3), Thermo Fisher) was added to a final concentration of 950 nM, and cells were filtered in 5 ml cell strainer tubes (Falcon). DiBAC4(3) was diluted every day to exchange buffers from DMSO to Tyrode^44^. Cells were kept on ice and protected from light in the transfer to the flow cytometry core.

Each sample was run in entirety on a BD Fortessa H0081 flow cytometer. DiBAC4(3) was excited at 488 nm and recorded at 525/50 bandpass, and miRFP670 fluorescence was excited at 640 nm and recorded with a 670/30 nm bandpass filter. Cells were gated on Side Scattering and Forward Scattering to separate out whole HEK293FT cells, gated on forward scattering area and width to separate single cells, then gated on co-expressed miRFP670 to gate out cells that received a plasmid, then a gate was set on the lower 50% of a histogram of wildtype Kir2.1 function, all mutants percentage of cells in this gate are reported. The analysis was performed in FlowJo 10 (FlowJo, LLC).

#### Flow Cytometry Assay for Light-modulation of Kir2.1 function

The generation of all single mutants used in the optogenetic switching assay was previously described in the resting membrane potential assay methods section. A Cry2-P2A-mKate2 domain was generated using gene fragments (Gen9) and assembled into the expression vector pEGFPN3 (Invitrogen) using BsmbI-mediated Golden Gate cloning ^63^. The Kir2.1(V302M) mutant was generated using Q5 site-directed mutagenesis (NEB).

The light-modulation assay was conducted for Cib81 mutants chosen as representative examples for the various permissibility and functional phenotypes we had observed. In addition, negative controls such as wildtype Kir2.1 and a pore dead mutant V302M ^45^ were included. 4 ug of each mutant, 3ug of dummy plasmid (pATT-Dest), and 100 ug of Cry2-P2A-mKate2 were transiently transfected using 6ul PEI across 16 wells of a 24 well dish at 20% confluency. Cells from each well were detached using 300 ml accutase, washed three times, and resuspended in 4ml Tyrode. DIBAC4(3) was added to a final concentration of 950 nM, and cells were filtered in cell strainer 5 ml tubes (Falcon). Filtered cells were divided into twelve 5 ml tubes (300ul each) and kept on ice and protected from light in the transfer to the flow cytometry core.

Cells expressing each mutant, wt Kir2.1, Kir2.1(V302M) were challenged by the addition of K-gluconate at different concentrations (5 mM, 10 mM, 15 mM, 25 mM, 40 mM, and 70 mM), with and without illumination (455 nm LED (Thorlabs), 30 seconds, 100% duty cycle, 100uW/mm^2^).

Each sample was run in entirety on a BD Fortessa H0081 flow cytometer. DIBAC4(3) was excited at 488 nm and recorded with 525/50 bandpass and 502 long pass filters, miRFP670 was excited at 640 nm and recorded with a 670/30 bandpass filter, and mKate2 was excited at 561 nm and recorded at 610/20 bandpass and 595 long pass filters.

Each sample was recorded for 5 minutes or until completion. Cells were gated on Side Scattering and Forward Scattering to separate out whole HEK293FT cells, gated on forward scattering area and width to separate single cells, then gated on co-expressed miRFP670 (Kir2.1 mutant) to gate out cells that received a mutant plasmid. For each paired sample (dark and light) a custom gate was created in the non-illuminated sample to include the 15% most hyperpolarized cells (using the flowStats package^69^). The number of events falling into this gate were then compared to the corresponding illuminated sample using the Chi-Squared test and reported as Dissimilarity (X^2^, light vs. dark). Dissimilarities at different K^+^ challenges were normalized to correct for photobleaching and averaged.

#### Patch Clamp Electrophysiology

HEK293FT cells were transiently transfected with Kir2.1 (wt) or Kir2.1 insertion mutant and Cry2-P2A-mKate using PEI. Cells were screened for mKate2 expression using a 565nm high-power LED (Thorlabs) filtered by a 560±40nm bandpass filter (Semrock) through a 40X lens. K^+^ currents were recorded 36-48 hours post-transfection using on-cell patch clamp electrophysiology. Patches with clear channel activity were stimulated with blue (455nm) light delivered by a LED (Thorlabs) at 100 uW/mm^2^ for 50 seconds at 100% duty cycle. Analog signals were filtered (2-5 kHz) using the built-in 4-pole Bessel filter of a Sutter Instrument IPA patch clamp amplifier, digitized and stored. Bath solution contained 125mM NaCl, 2mM KCl, 3mM CaCl_2_, 1mM MgCl_2_, 10mM HEPES, 30mM glucose, adjusted to pH 7.3 with NaOH. The pipette solution contained: 125mM K-Gluconate, 8mM NaCl, 0.1mM CaCl_2_, 0.6mM MgCl_2_, 1mM EGTA, 10mM HEPES, 4mM Mg-ATP, 0.4mM Na-GTP, adjusted to pH 7.3 with KOH. Osmolarity was adjusted to 295 - 300 mOsm with sucrose. Electrodes were drawn from borosilicate patch glass (Warner Instruments) to a resistance of 2-6 MΩ.

Data analysis was done using custom R scripts. After correction for baseline drift, representative all points current amplitude histograms for sweeps (1) before, (2) during, and (3) after > 2 min after illumination were calculated. Histograms were fit to a sum of Gaussian functions (n = 2–6):

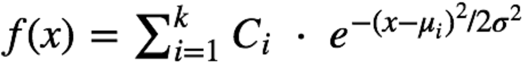

Where *f(x)* is the count of points in a bin with current amplitude *x*, with the mean μ, standard deviation σ, and *C* total number of points belonging to the *i*th component. In a case with a single channel, the open probability is calculated as:

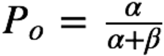

with α and β, the total number of points belonging to the Gaussians that describe the open and closed state, respectively.

---

**Figure S1:**
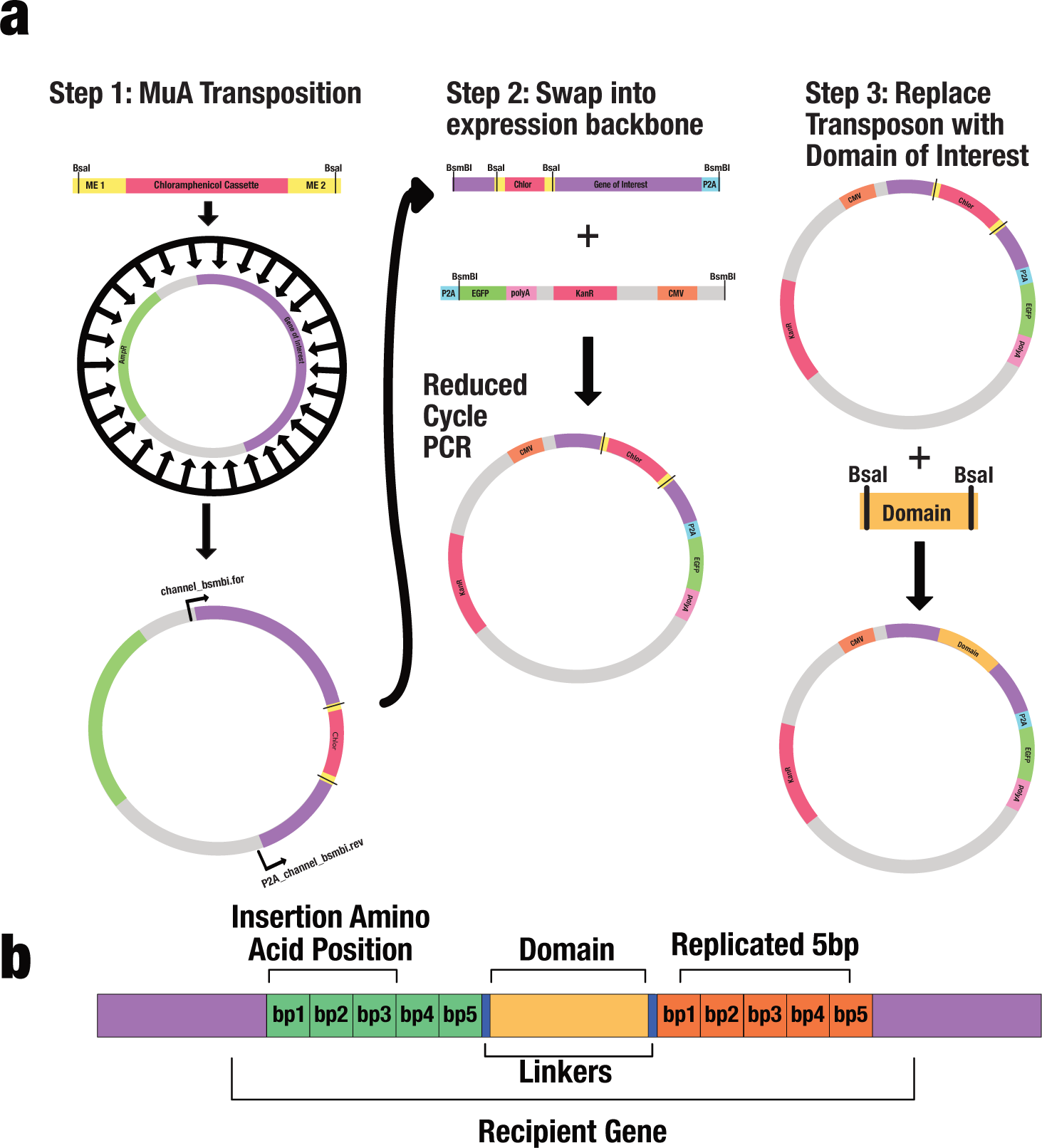
Insertion Library Construction. **(a)** Libraries were generated in three cloning and selection steps. (1) Select for a MuA transposase delivered engineered MuA transposon with a chloramphenicol antibiotic cassette in a plasmid carrying Kir2.1. Flanking the antibiotic cassette are the beginnings and ends of flexible linkers with golden gate compatible BsaI type IIS restriction sites. (2) Reduced cycle PCR amplify and add on golden gate compatible BsmBI type IIS restriction sites and size separate channel genes with inserted transposons from those without transposons. Insert channel gene into a mammalian expression vector in-frame with a P2A-EGFP cassette. (3) Replace the transposon with a PCR amplified domain of interest with complementary BsaI sites and linkers using BsaI mediated golden gate cloning. **(b)** Architecture of a domain insertion position: At the position of the domain insertion the five positions upstream are replicated on the other side of the transpositions; domain insertion positions are identified as the last full codon coding for an amino acid or in other words that corresponding to the amino acid coded by bp1-3 of the replicated sequence. Domains are inserted with linkers to bring it into frame after insertion at +2 reading frame relative to the coding sequence.

**Figure S2:**
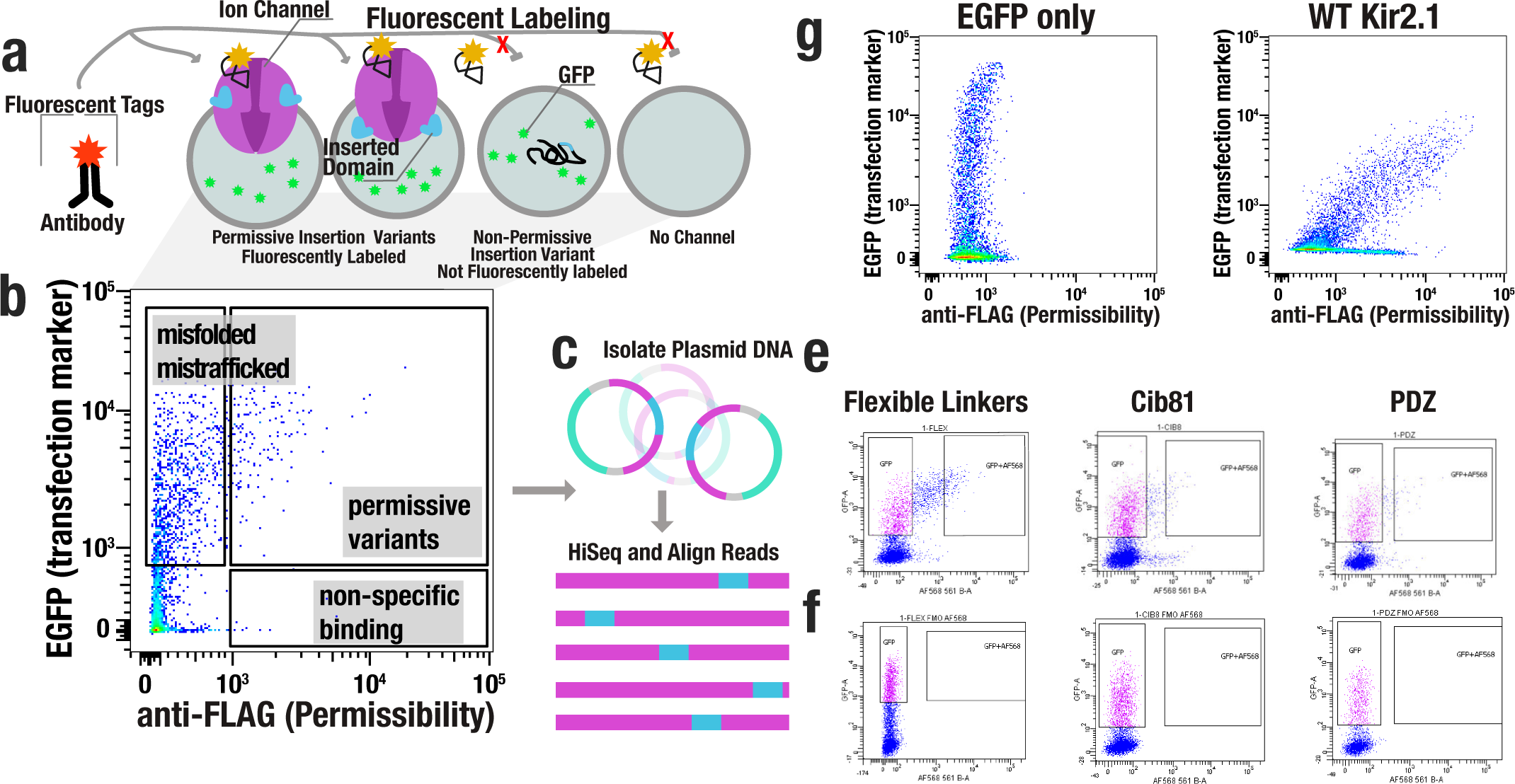
Permissibility Assay. **(a)** Express domain insertion libraries in HEK293 cells and label with anti-FLAG Alexa568. (**b**) Use Flow Sorting to isolate cells into two populations: surface expressed (permissive insertion variants) and non-surface expressed (non-permissive) insertion variants. (**c**) Isolate plasmid DNA from each population and subject to HiSeq. (**d**) Labeling controls (only GFP and wildtype Kir2.1-p2a-EGFP) were expressed in HEK293 to demonstrate antibody labeling. (**e**) Examples for each anti-FLAG labeled sorted samples (GSAGx2 and GSAGx3, Cib81 and PDZ) expressed in HEK293 cells with gates that were used for sorting. (**f**) Identical samples as in (**e**) without anti-FLAG labeling to guide setting gates.

**Figure S3:**
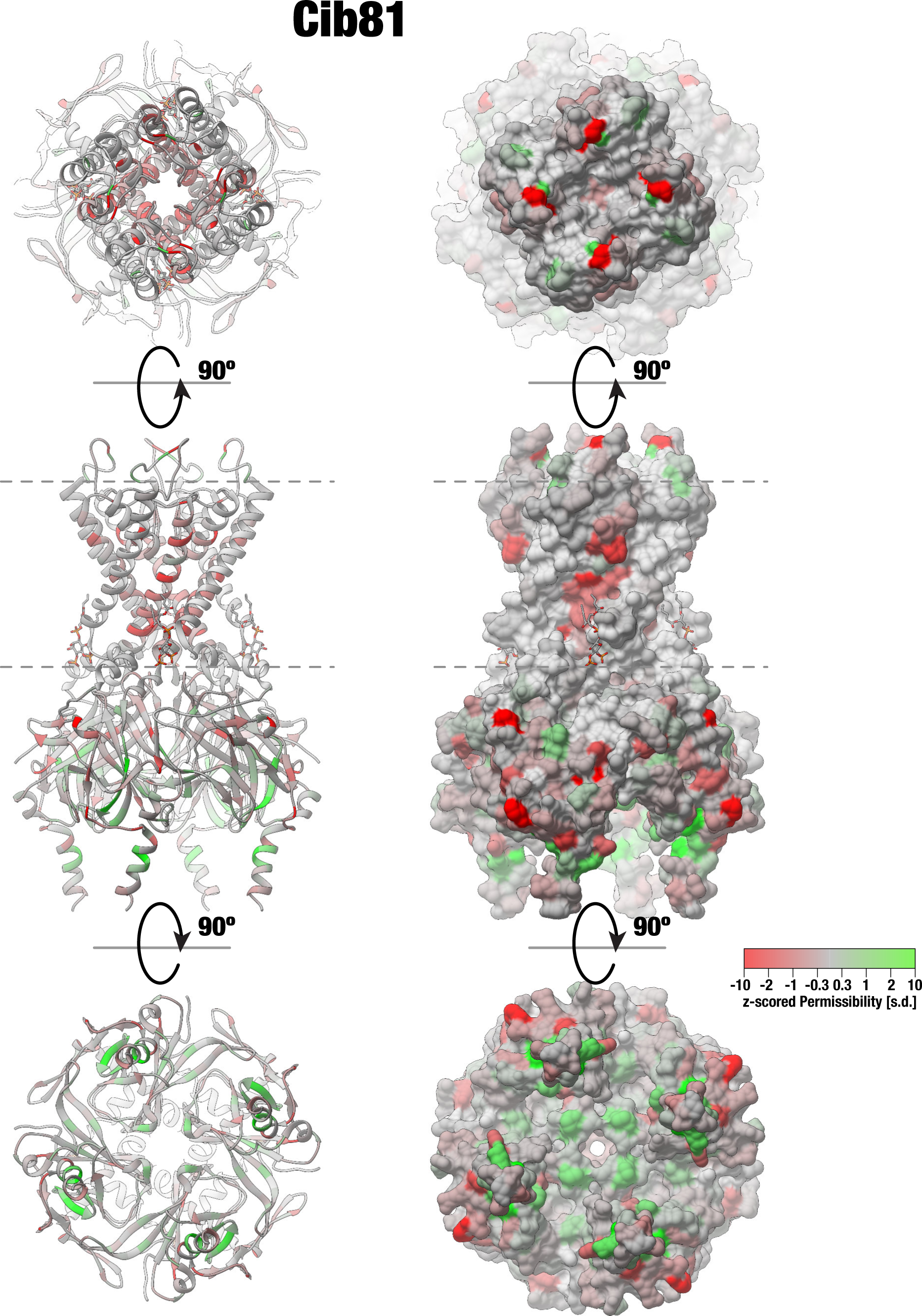
Domain Insertion Permissibility. Permissibility data for inserting Cib81 is mapped on the crystal structure of chicken Kir2.2 (PDB 3SPI) displayed as a ribbon (left) or surface model (right). Dashed lines indicate plasma membrane boundaries.

**Figure S4:**
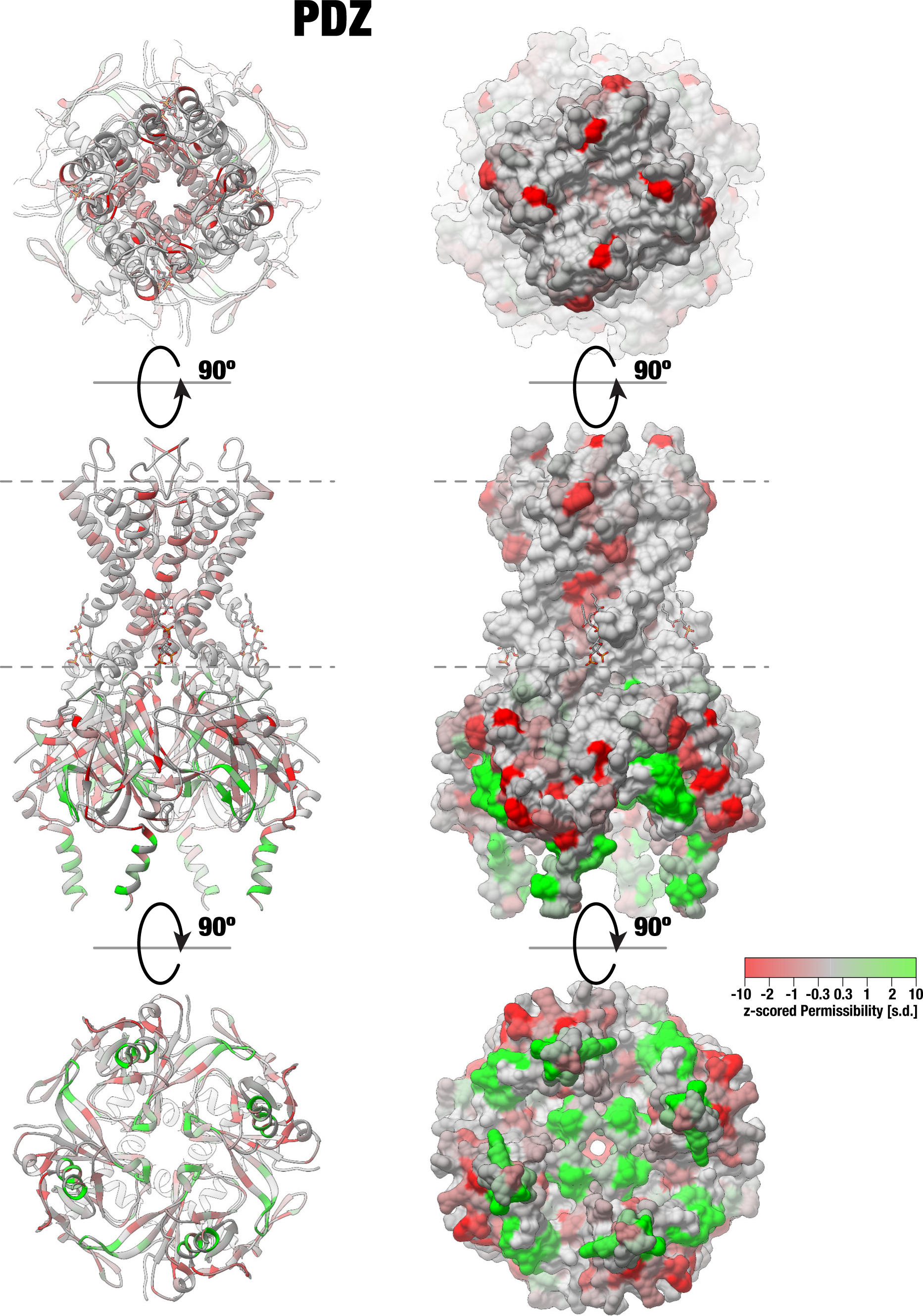
Domain Insertion Permissibility. Permissibility data for inserting PDZ is mapped on the crystal structure of chicken Kir2.2 (PDB 3SPI) displayed as a ribbon (left) or surface model (right). Dashed lines indicate plasma membrane boundaries.

**Figure S5:**
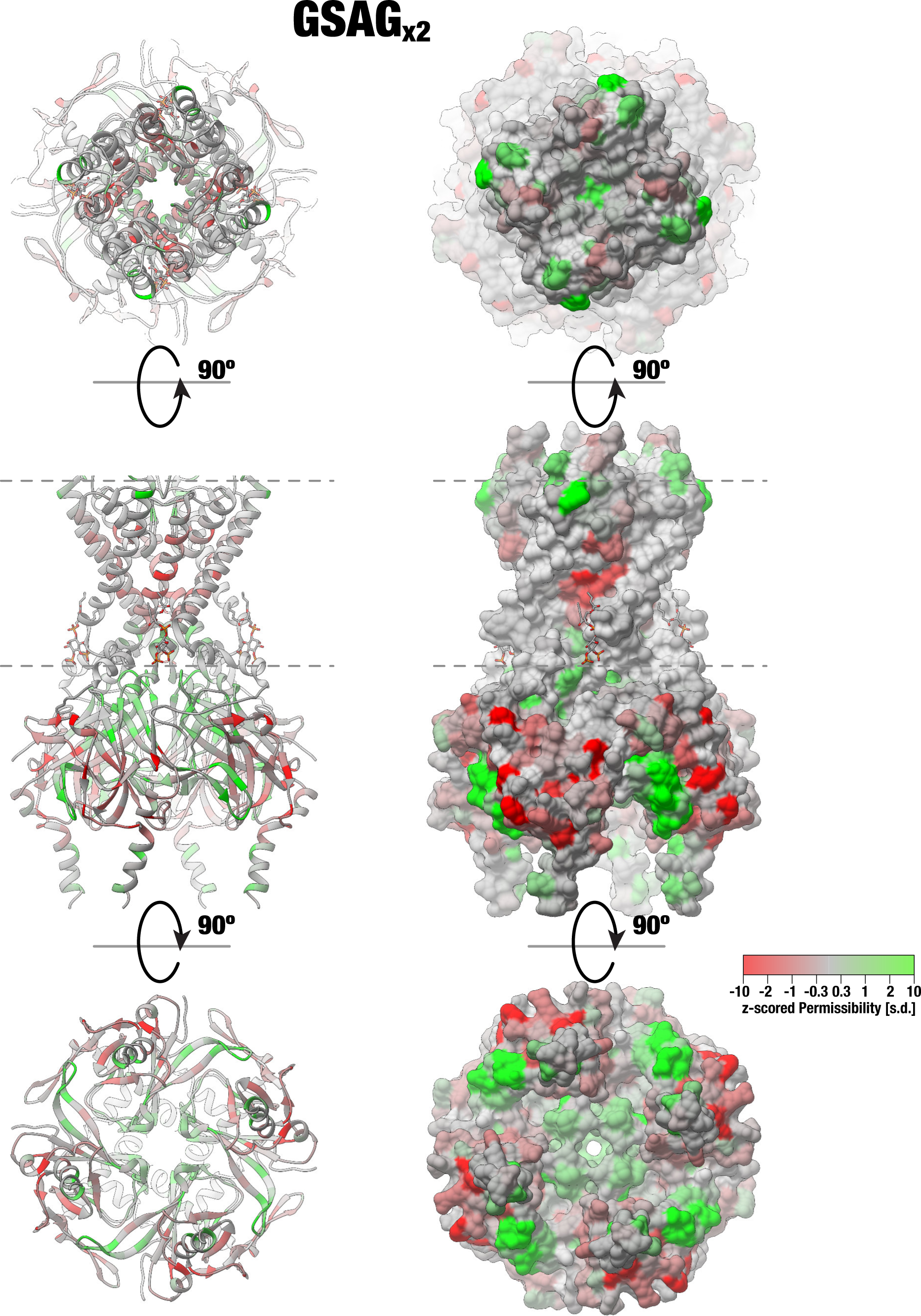
Domain Insertion Permissibility. Permissibility data for inserting GSAG_x2_ linkers is mapped on the crystal structure of chicken Kir2.2 (PDB 3SPI) displayed as a ribbon (left) or surface model (right). Dashed lines indicate plasma membrane boundaries.

**Figure S6:**
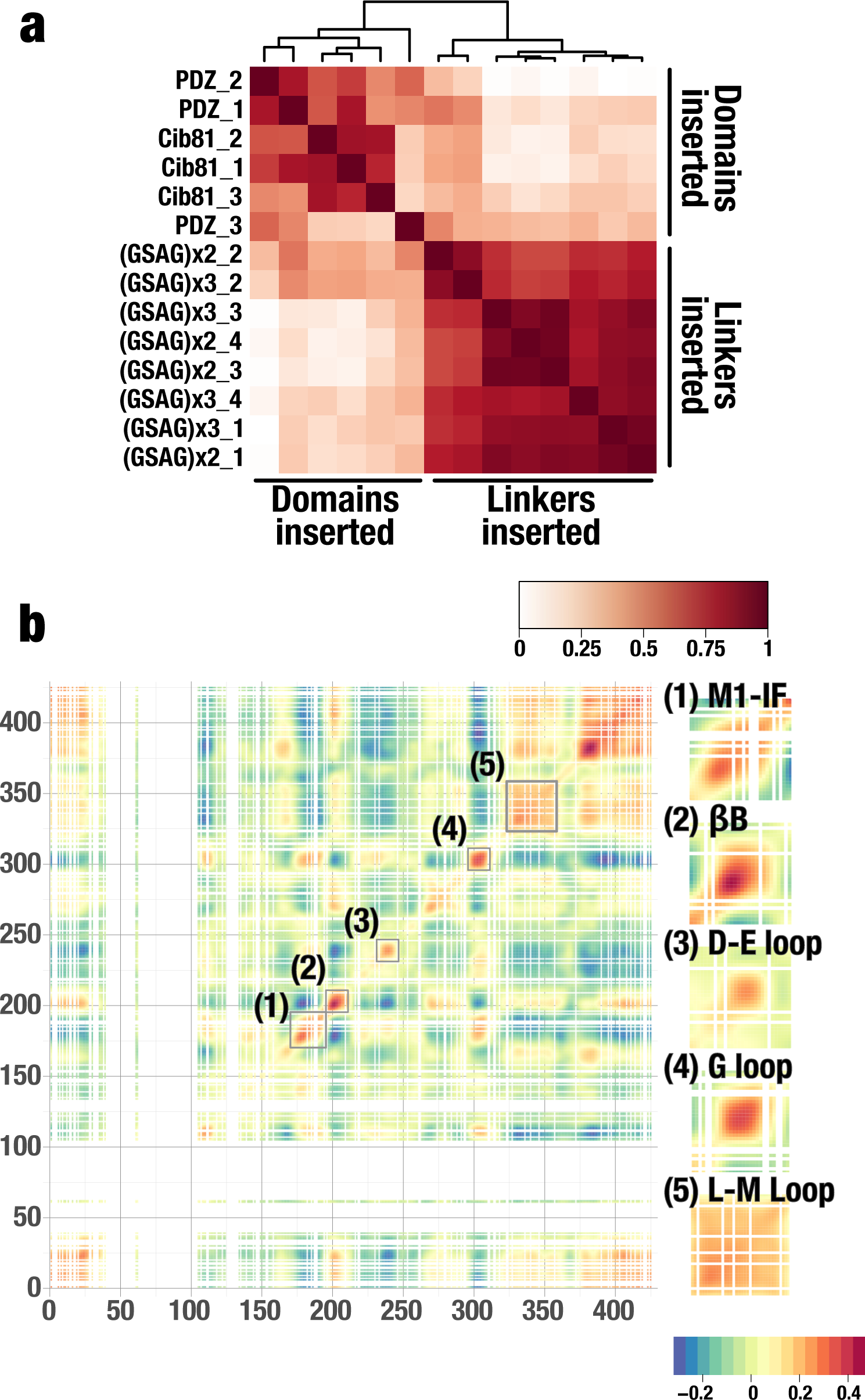
Pearson correlation of biological replicates and site-specific permissibility correlations. **(a)** Hierarchical clustering of all individual permissibility datasets based on domain structure and length. Biological replicates show a high degree of reproducibility. As expected, datasets for structured domains (Cib81 and PDZ) cluster discretely with themselves than with each other whereas flexible linkers cluster indiscriminately between each other. **(b)** In Kir2.1, permissibility is highly correlated in (1) M1-IF – the PIP_2_ binding site, (2) the βB loop where ATP binds in Kir6.2, (3 & 5) the βD-βE and βL-βM loops where Gβγ binds in GIRK, as well as (4) the G loop involved in channel gating. White indicated sites with incomplete data.

**Figure S7:**
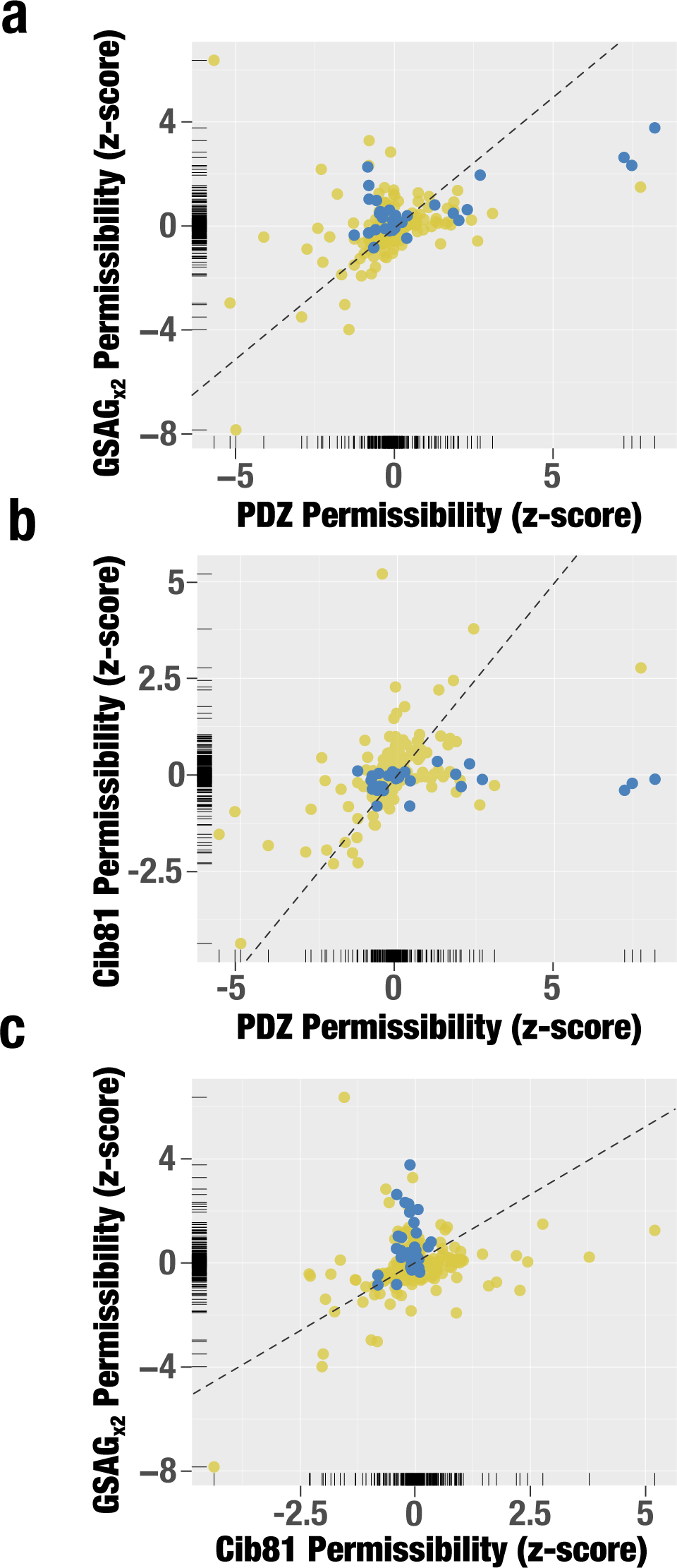
Differential Permissibility is common in functionally important sites. Z-scored permissibility comparisons between PDZ and GSAG_x2_ linkers (**a**), PDZ and Cib81 (**b**), and Cib81 and GSAG_x2_ linkers (**c**). Each dot represents a site in Kir2.1, color-coded by function (yellow, not involved in allosteric regulation; blue, involved in allosteric regulation). Equivalent permissibility values for both insertion type are indicated by the dashed line.

**Figure S8:**
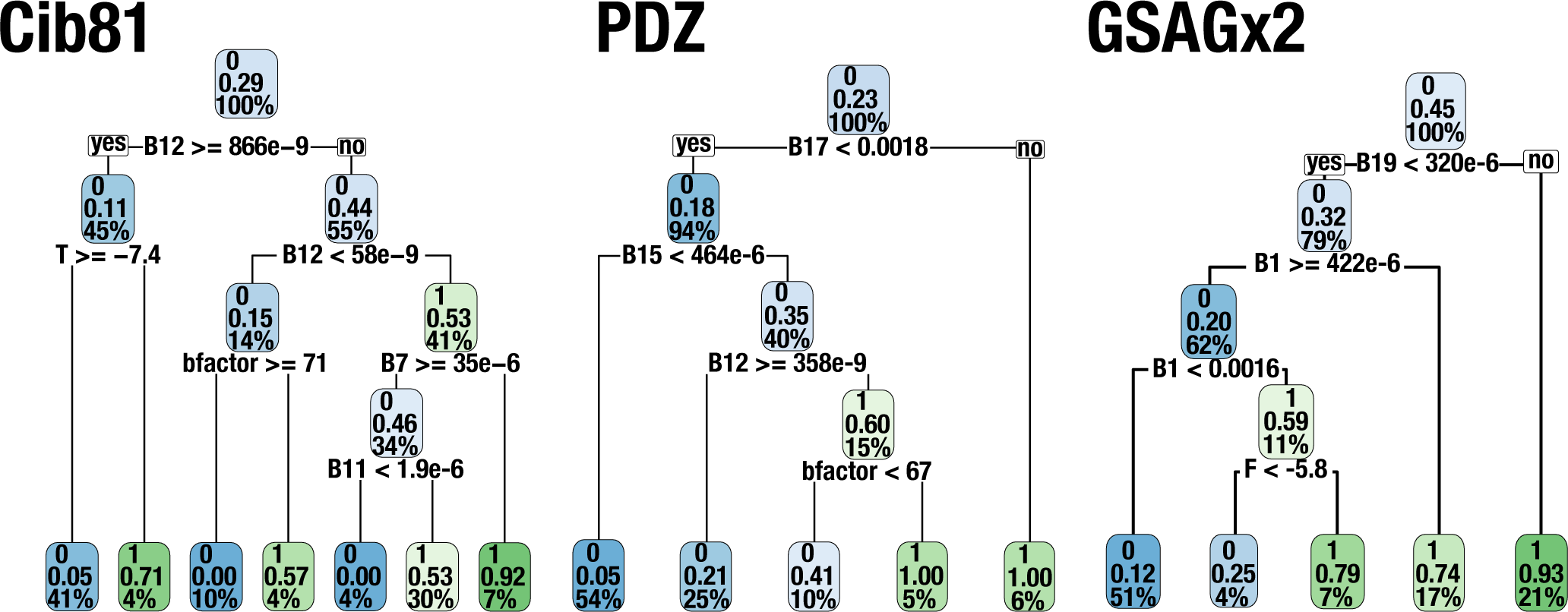
Decision Trees. Decision trees that were trained on calculated conservation, static structural and dynamic protein properties to predict binarized Cib81, PDZ and GSAG_x2_ permissibility. Models overwhelmingly use dynamic properties (normal modes) over all other categories of computed properties. All trees were restricted to a maximum depth of four and cross-validated ten times to minimize overfitting and enable comparison between models. Decision tree leaves can be read as: the topmost number refers to the leaf class (0-not permissible and 1-permissible), percentage of permissible samples within the leaf that fall within the class, and percentage of all data that are within the leaf. Color intensity refers to how pure the leaf is not permissive (blue) or permissive (green).

**Figure S9:**
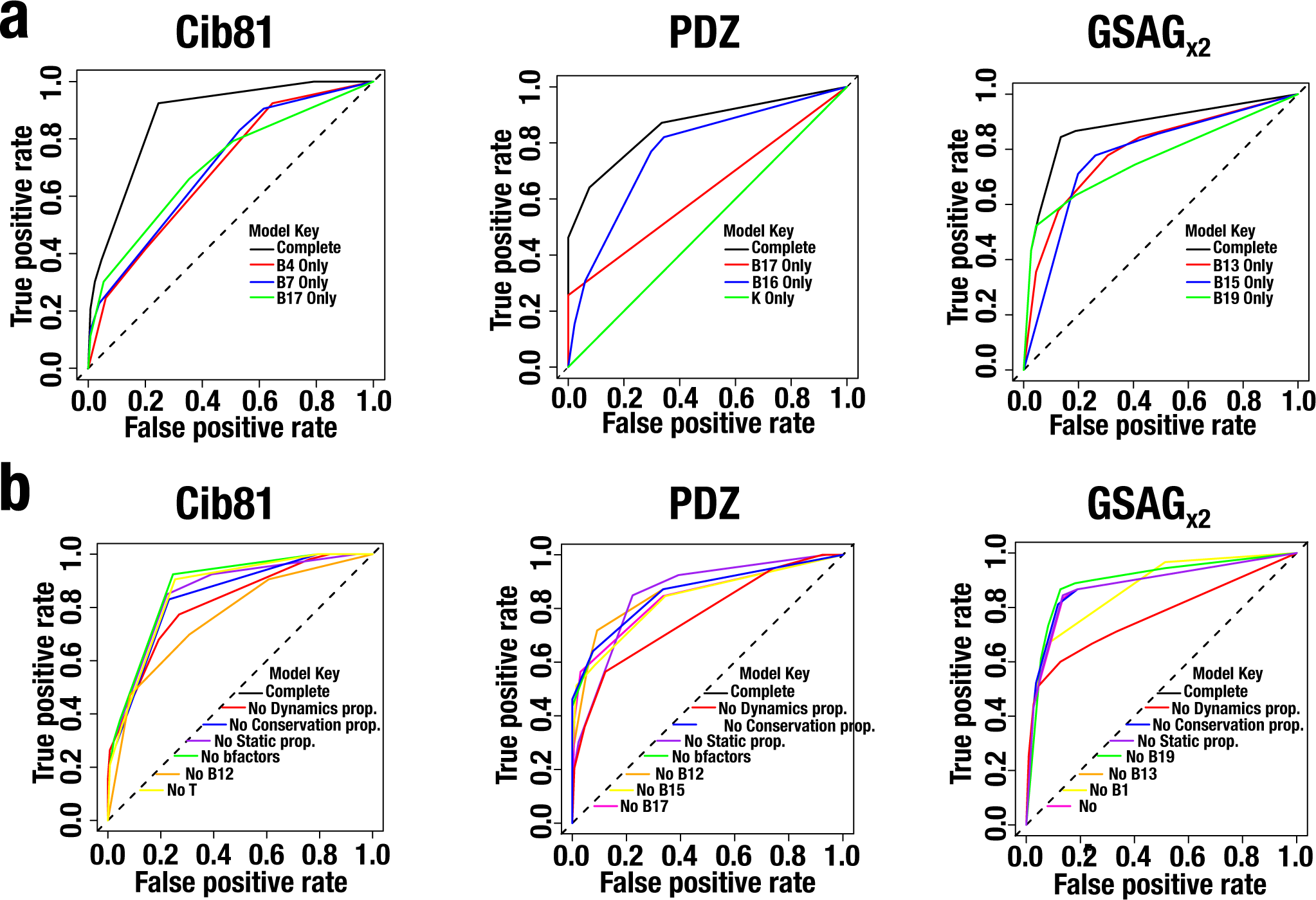
Decision trees trained on limited numbers of properties. **(a)** Decision trees were trained using the top three Spearman co-efficient correlated properties individually using the same model parameters as before (a maximum depth of four and cross-validated ten times). Models’ performance was compared using receiver operator characteristic (ROC) curves. All models perform worse than models able to train on single properties showing that multiple interacting properties determine permissibility. Interestingly, the predicted effect of any amino acid mutated to lysine was highly linearly correlated with PDZ permissibility, however, using it alone was not able to predict PDZ permissibility at all whereas normal modes were able to predict PDZ permissibility individually. **(b)** Decision trees were trained with the decision tree determined predictive properties withheld individually and withholding each entire class of properties (conservation-, static structural- and dynamics based properties) using the same model parameters as before (a maximum depth of four and cross-validated ten times). Models’ performance was compared using ROC curves. In every case, removing all computed conservation and static structural properties had little effect on model performance, however, removing dynamics based properties was detrimental to permissibility predictions. Apart from two examples (Cib81~B12 and GSAG_x2_~B1) removing individual properties were not substantially impactful on model performance meaning there is redundancy in computed properties. Overall, all these decision trees reinforce that (1) dynamics are the most informative and closest correlated protein properties to permissibility and (2) permissibility is non-linearly made up of multiple interacting protein properties.

**Figure S10:**
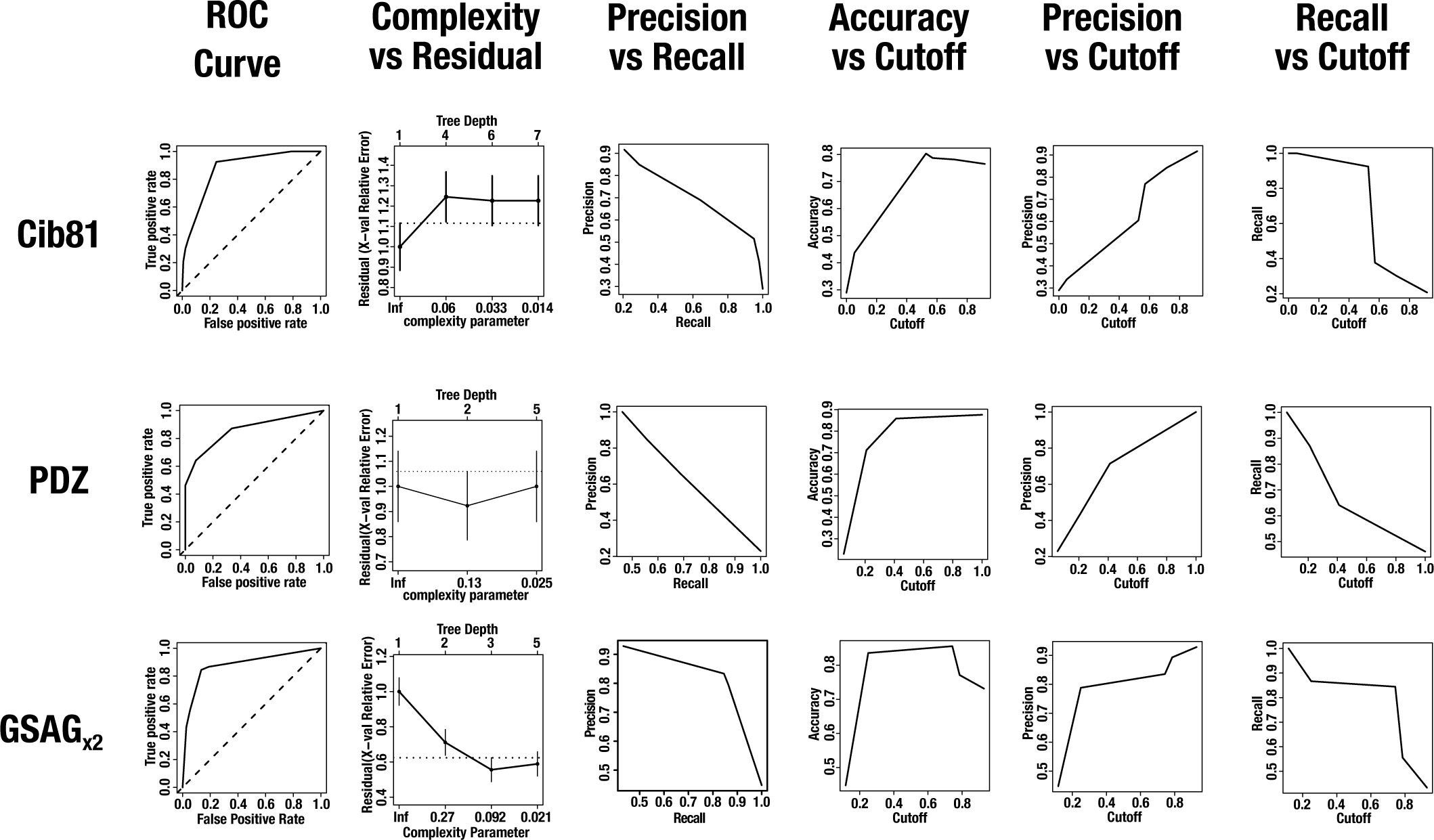
Decision Tree Model Performance. Cib81, PDZ, and GSAG_x2_ decision tree performance using different criteria. All models based on all parameters perform far better than random.

